# Single cell RNA analysis uncovers the cell differentiation trajectories and functionalization for air breathing of frog lung

**DOI:** 10.1101/2023.07.18.549571

**Authors:** Liming Chang, Qiheng Chen, Bin Wang, Jiongyu Liu, Meihua Zhang, Wei Zhu, Jianping Jiang

**Author notes:** **Corresponding authors:** Jianping Jiang,; Wei Zhu.

## Abstract

The evolution and development of vertebrate lungs have received extensive concerns for the significance in terrestrial adaptation. Amphibians possess the most primitive lungs among tetrapods, underscoring the evolutionary importance of lungs in bridging the transition from aquatic to terrestrial life. However, the intricate process of cell differentiation during amphibian lung development is still lacking. Using single cell RNA-seq, we identified 21 cell types in the developing lung of a land-dwelling frog (*Microhyla fissipes*). We elucidated that single type of alveolar epithelial cells (AECs) existed in amphibian and the diversity of AECs may correspond to the capacity for terrestrial adaptation in tetrapods. Based on pseudotime trajectories analysis, we revealed previously unrecognized developmental-specific transition cell states of epithelial and endothelial cells supporting the rapid morphogenesis of lung during metamorphic climax. We illustrated the cellular and molecular processes during lung functionalization. These findings uncover the cell differentiation trajectories and functionalization for air breathing of frog lungs and highlight the evolutionary peculiarity of the primitive lungs.

**Graphical abstract:** 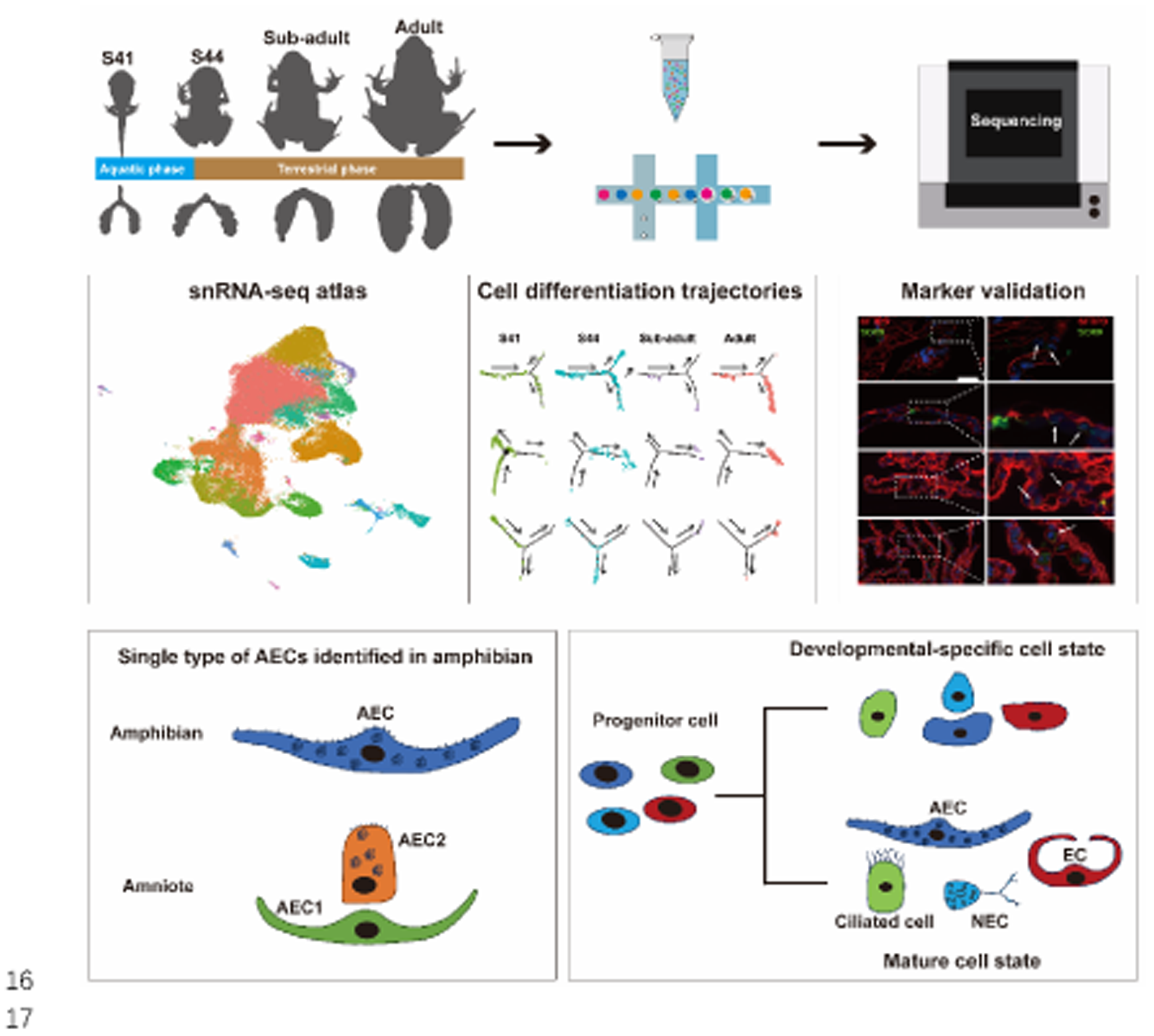

## Introduction

The water-to-land transition is a milestone in the evolution history of vertebrates. The emergence of air-breathing organ corresponds with the shift of oxygen medium from water to air and facilitates vertebrate terrestrialization (Hsia et al., 2013; Pike, 1924). In tetrapods, lungs serve as the air-breathing organ, and its functionalization is a tightly regulated multistage process that involves the ventilation, oxygenation, and further perfusion of the pulmonary microcirculation (Burggren and Infantino, 2015; Morton and Brodsky, 2016; Zoetis and Hurtt, 2003). To perform the highly choreographed function, 4 major types of cells play important roles during lung development, i.e., epithelial cells, endothelial cells (ECs), mesenchymal cells (MCs), and pulmonary circulating blood cells. In addition, there are more than 20 types of cells in terms of these 4 major cell types (Guo et al., 2019). With the development of high-throughput single cell RNA sequencing (scRNA-seq) technology, a large number of pulmonary single cell atlases have been constructed for various organisms, such as *Homo sapiens* (He et al., 2022; Kadur et al., 2022), *Mus musculus* (Cui et al., 2019), non-model mammals, reptiles and birds (Chen et al., 2021), and *Xenopus laevis* (Liao et al., 2022). These proceedings have been providing deep understandings of the pulmonary cell differentiation across the vertebrate groups from water- to land-dwelling. Therefore, cell differentiation of vertebrate lungs in evolutionary studies becomes a hotspot. However, the dynamic processes of pulmonary cell differentiation and functionalization during vertebrate lung development are largely lost and relatively less concerned, particularly for the water-to-land transitional groups.

Alveolar epithelial cells (AECs) are essential components of the blood-gas barrier in the lung and play a vital role in gas exchange. Ultrastructure of AECs were studied in amniotes, including snakes (Maina, 1989), lizards (Klemm et al., 1979), turtles (Fleetwood and Munnell, 1996), birds (Maina and King, 1989) and mammals (Jansing et al., 2017; Metzger et al., 2008). It is widely supported that there are two distinct types of AECs in amniotes. Single-cell atlas of the lung were constructed for 11 non-model species of reptiles, birds, and mammals, further confirming that there were two types of AECs in amniotes (Chen *et al*., 2021). Amphibians, as a crucial vertebrate group that achieved the transition from aquatic to terrestrial life, possess the most primitive lungs among tetrapods (Okada et al., 1962; Perry and Sander, 2004). The presence of two types of AECs in amphibians was initially described by Okada et al. (1962) based on the relative distance of cell bodies to capillaries. Goniakowska-Witalińska (1978, 1986) proposed that there was only one type of AECs, as AECs do not exhibit significant ultrastructural differences in amphibian (Goniakowska-Witalińska, 1978; 1986). Rankin et al. (2015) observed one type of AECs in *X. laevis* base on in situ hybridization studies. They also proposed the possibility of different subtypes of AECs that may not have been identified as the limitations of technology (Rankin et al., 2015). Recent, only one type of AECs was identified with the pulmonary single cell atlases in *X. laevis* that exclusively inhabit aquatic environments throughout their life cycles (Liao *et al*., 2022). Whereas, the differentiation of AECs in amphibians is still unclear and further scRNA sequencing studies on AECs are needed particularly in species with distinct aquatic and terrestrial life stages.

Most amphibians undergo metamorphosis to adapt for the terrestrial environment with the transition of respiratory organs from gills to lungs (Burggren and Infantino, 2015). During amphibian metamorphosis, the rapid morphogenesis and functionalization of lungs involved dynamic structural, biochemical, and physiological changes (Chang et al., 2022). All these processes are depended on the cellular proliferation, differentiation, and interaction of multiple types of cells. Moreover, the lung has low rates of cell turnover in adult frog, making it challenging to capture developmental-specific transition cells states (Blenkinsopp, 1967; Rawlins and Hogan, 2008). A high-resolution cell atlas of lung during amphibian metamorphosis will identify developmental precursors and transition cell states and predict cell differentiation trajectories. It will be conducive to a better understanding of lung functionalization for air breathing.

*Microhyla fissipes* (Anura: Microhylidae) is a good model for exploring lung functionalization for air breathing during the transition from water to land, as their life cycles comprise of distinct aquatic and terrestrial phases (Liu et al., 2016; Wang et al., 2017). Our previous studies have been addressed the morphological changes and molecular mechanism of lung development by using Micro-CT and whole tissue sequencing during *M. fissipes* metamorphosis (Chang *et al*., 2022), while the function of individual cells during lung development are not clear. In this study, scRNA-seq analysis were applied to further elucidate three key issues: (1) the classification of AEC types in amphibians; (2) the differentiation trajectories and transition cell states of epithelial cells and ECs; (3) the cellular and molecular processes of lung morphogenesis and functionalization during frog metamorphosis. This study gives a systematic view on the dynamic variations of pulmonary cell differentiation and functionalization for air breathing of frog lungs and shed some light on the evolutionary peculiarity of the primitive lungs.

## Result

### scRNA-seq sequencing and unbiased clustering of developmental lung cells

Single cells of *M. fissipes* lungs were obtained from 8 samples of 4 developmental stages, namely S41, S44, Sub-adult (post metamorphosis), and Adult, representative of the aquatic stage, early, middle, and late stages of terrestrial transition, respectively (Figure 1A). Having passed quality control filtering (Supplementary Data 1), a total of 69,490 lung cells were performed unbiased clustering, and these cells can be grouped into 4 major types including pulmonary circulating blood cell, MCs, epithelial cell, and ECs (Figure 1B), which can further be divided into 21 cell clusters (Figure 1C; Supplementary Figure 1A) in an unsupervised manner (k-means method) using uniform manifold approximation and projection (UMAP).

**Figure 1.**
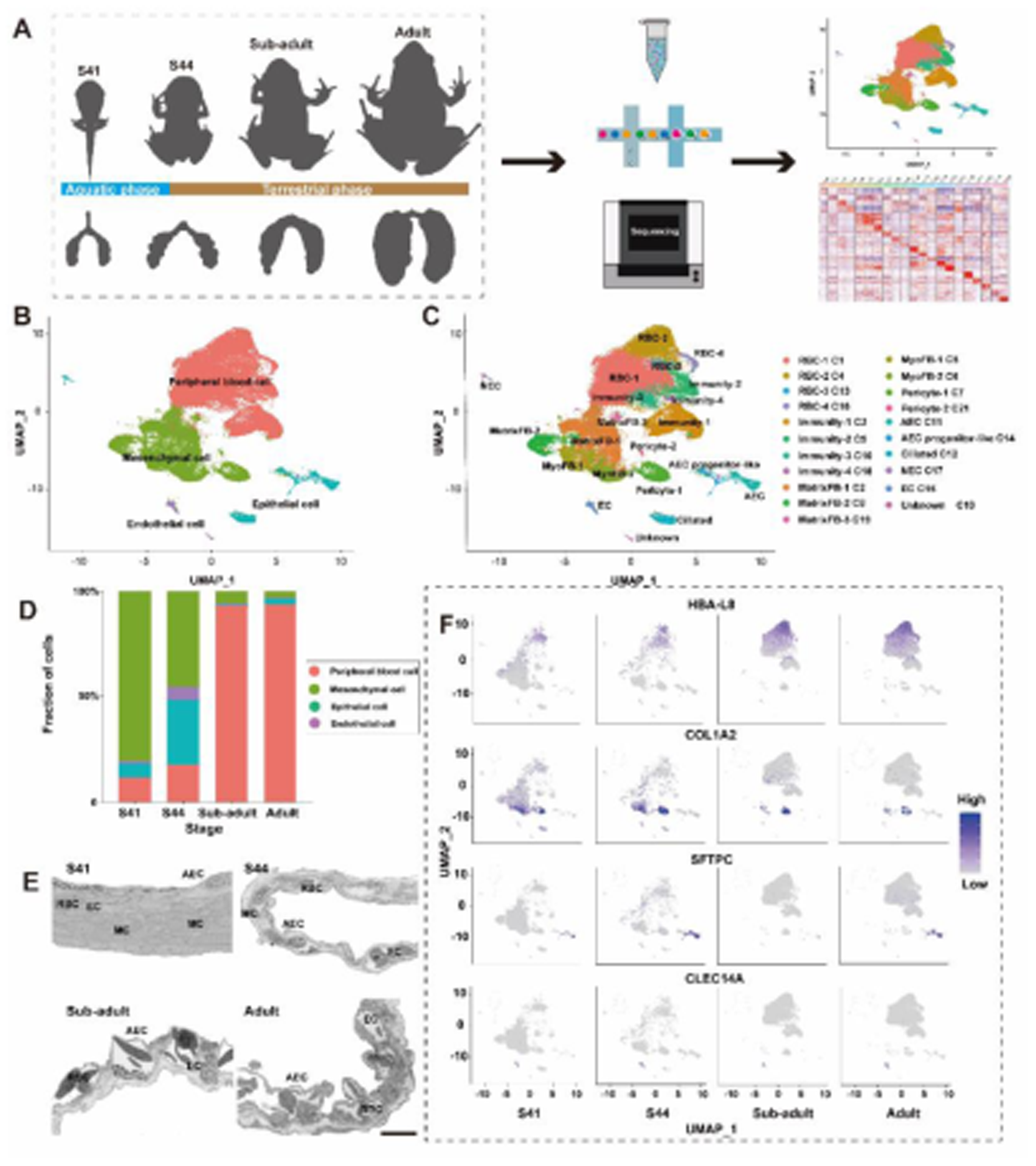
Cellular diversity of lungs in 4 developmental stages. (A) A schematic of the basic workflow for the pulmonary cell landscape in *M. fissipes* using the 10× Genomics platform. (B) Uniform manifold approximation and projection (UMAP) plot showing four major cell types in lung. (C) UMAP plot showing 21 cell types in lung. (D) Relative proportion of each major cell types. (E) Transcriptional dynamics of marker genes showing the proportional variations in the 4 major cell types. AEC, alveolar epithelial cell; EC, endothelial cell; MC, mesenchymal cell; RBC, red blood cell. (F) Ultrastructural dynamics of alveoli showing the proportional variations in the four major cell types, scale bar, 10 *μ*m.

Cell clusters were annotated based on the expression of conservative cell markers from Cell Marker 2.0 Database (Hu et al., 2022) and differential expression genes (DEGs) profile of each cell cluster (Supplementary Figure 1A; Supplementary Data 2). In brief, Cluster 1 (C1), C4, C13, and C16 were identified as red blood cells (RBCs) due to their high expression levels of *HBA-L8*; C15 was classified as endothelial cells (ECs), distinguished by its specific expression of *CLEC14A* (Dusart et al., 2019); C11 was categorized as AECs, characterized by the specific expression of *SFTPB* and *SFTPC* (Hurley et al., 2020; Travaglini et al., 2020; Xu et al., 2016); the MCs comprised seven clusters including C2, C5, C6, C7, C8, C20, and C21, which exhibited high expression levels of MC marker genes such as *COL1A1* and *COL1A2* (Guo *et al*., 2019); C17 specifically expressed neuroendocrine cell (NEC) marker *ASCL1* (Bischoff et al., 2021; Plasschaert et al., 2018). Among the MCs, C5 and C6 were characterized by the highly expressed myofibroblast (MyoFB) marker *ACTA2* (Nakahara et al., 2021), while C7 and C21 specifically expressed the pericyte marker *RGS5* (Adams et al., 2020; Whitsett et al., 2019) (Supplementary Figure 1B). To assess the validity of the scRNA-seq data, we compared the proportions of cell types between samples. Importantly, no significant differences were observed in the cell proportions between the two biological replicates within the same group or stage (Wilcoxon rank-sum test, *p value* > 0.5, Supplementary Figure 1C). This finding strongly supports the high quality and reliability of the scRNA-seq data obtained in this study.

In regards to cell dynamics, C16 was not captured in S41, while C20 and C21 were not captured in sub-adult and adult, respectively. The rest of cell clusters were present in all of the 4 developmental stages. MCs were the predominant major cell type at S41 and S44 and then decreased in proportion with the proceeding of development. In contrast, the proportions of pulmonary circulating blood cells increased with development and accounted for the largest cell proportions at sub-adult and adult stages. Notably, the transition from S41 to S44, the initial terrestrial stage, was characterized by an opposite proportional variation in MCs and epithelial cells, and the latter one showed a drastic increasement (Figure 1D; Supplementary Figure 1D). The expression patterns of the respective marker genes (Figure 1E) and alveoli ultrastructure (Figure 1F) supported the proportional variations of the 4 major cell types in lung. These results suggested a quick cellular transitions during pulmonary morphogenesis.

### Diverse MCs during lung morphogenesis and functionalization

Seven types of MCs were identified in *M. fissipes* lungs, including three types of Matrix Fibroblasts (MatrixFBs), two types of MyoFBs, and two types of pericytes (Supplementary Figures 2A, B). All of the three MartrixFBs showed much higher abundance at larval stages (Figure 2A). The MatrixFB-1 and MatrixFB-3 differ in the expression level of MartrixFB marker genes (Figure 2B), but converge in the cellular functions, e.g., collagen-containing extracellular matrix, angiogenesis, and osteogenesis (Supplementary Figures 2C-E). The transcriptional factors (TFs) of these two MartrixFBs (MartrixFB-1: *FBN2*, *FOXF1-B*, *NOTCH2*, and *AEBP1*; MatrixFB3: *HAND2*, *FLNC*, *BNC1*, and *ZEB2*) are responsible for regulating the multi-functionalization of MartrixFBs (Figure 2C). MatrixFB-3 seems to be a larvae-specific MatrixFBs for its absent in sub-adult and adult lungs (Figure 2A), and this is supported by the expression of *HOXC5*, a key regulator of cell positioning in early embryos development (Fritz and De Robertis, 1988). Unlike MatrixFB-1 and MatrixFB-3, MatrixFB-2 is featured by the genes involving in cell mitosis, chromatin organization (Figure 2B), and TFs regulating cell cycle (Figure 2C). And the expression of *HMGB2* and *HMGB3*, two critical regulators that balance self-renewal and differentiation in some stem cells (Abraham et al., 2013; Nemeth et al., 2006), suggests the self-renew potential of MatrixFB-2. These results suggest that different types of MatrixFBs in lung have a clear division of labor in self-renewal and functionalization.

**Figure 2.**
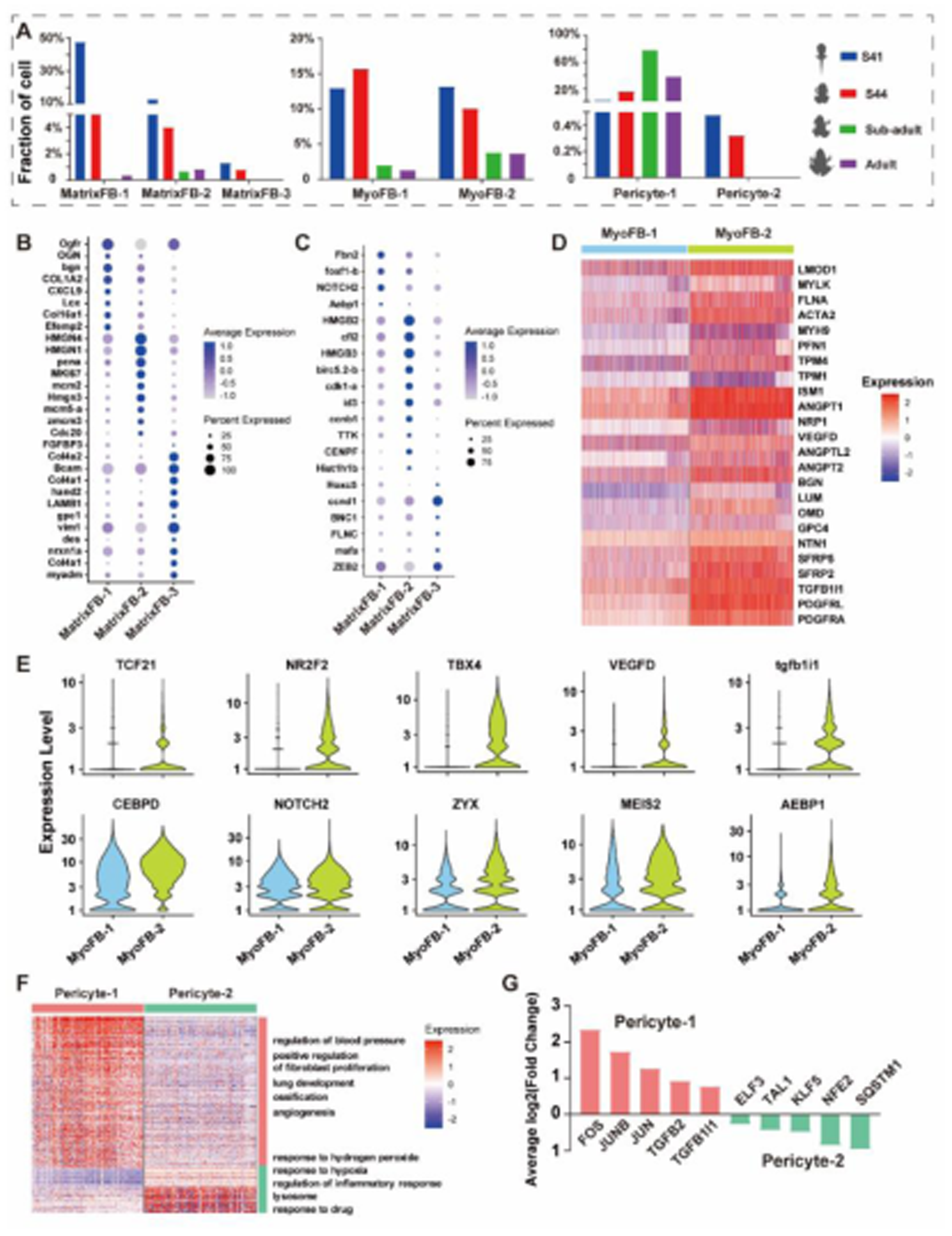
Diverse MCs and the molecular features. (A) Relative proportions of pulmonary MCs in four developmental stages. (B-C) Differential expression of genes (B) and TFs (C) among MatrixFBs. (D-E) Differential expression of genes (D) and TFs (E) among MatrixFBs between MyoFBs. (F) GO enrichment of DEGs between pericyte-1 and pericyte-2, adjusted *p value* < 0.001. (G) Differential expression of TFs between pericyte-1 and pericyte-2.

The two MyoFBs (MyoFB-1 and MyoFB-2) are characterized by the expression of *ACTA2* (Supplementary Figure 1B), and they occupy higher cell proportions during the larval stages than in juvenile and adult stages (Figure 2A). MyoFB-2 exhibits more robust expression of genes involved in collagen-containing extracellular matrix, myogenesis, and angiogenesis than MyoFB-1 (Figure 2D). In addition, MyoFB-2 extensively expresses *SFRP2* and *SFRP5*, which regulate cell growth, myogenesis and differentiation as modulators of WNT signaling (Deb et al., 2007).. Several TFs are highlighted in MyoFB-2 (Figure 2E), of which *TCF21* is involved in epithelial-mesenchymal interactions in lung morphogenesis (e.g., epithelial differentiation and branching morphogenesis) (Quaggin et al., 1999), *TBX4* has an essential role in the lung organogenesis (Zhang et al., 2013), *NR2F2* and *VEGFD* are active in angiogenesis ^36^. These results indicate a greater pluripotency of MyoFB-2, and we speculate that MyoFB-2 may be an active state of pulmonary MyoFBs which is essential for lung development.

Pericytes are mural cells of the microcirculation and wrap around the ECs that line the capillaries and venules (Birbrair et al., 2014; Dore-Duffy, 2008). Two types of pericytes were identified in *M. fissipes* lung (Supplementary Figure 2A, B). Pericyte-1 is more abundant in juvenile and adult stages, while pericyte-2 is specific to larvae stages (Figure 2A). Regarding to cellular functions, pericyte-1 is featured by angiogenesis and blood pressure regulation, while pericyte-2 is for immune processes and the maintenance of cellular redox balance. Pericyte-1 actively expresses 5 key TFs related to TGF-β receptor signaling pathway (i.e., *JUN*, *JUNB*, *FOS*, *TGFB2*, and *TGFB1I1*) (Figure 2G), which regulate angiogenesis and nervous system development (Gomes et al., 2005; Roberts et al., 1986). Pericyte-2 expresses *KLF5*, *ELF3*, and *SQSTM1*, which are responsible for angiogenesis (Guo et al., 2008), vascular inflammation (Rudders et al., 2001), and selective macroautophagy (Johansen, 2019), respectively.

### Epithelial cell heterogeneity and molecular function switches during metamorphosis

We identified 3 distinct types of epithelial cells, including AECs, ciliated cells, and NECs (Figure 3A). AECs are enriched for genes involved in epithelial cell differentiation, response to hypoxia, lamellar body, and microvillus (Figure 3A). Results of TEM indicated that AECs were filled by lamellar bodies and covered by dense microvilli (Figure 3B). The core functional proteins (e.g., *SFTPA1*, *SFTPB*, *SFTPC*, and *CAT*) are consistently expressed in AECs across the stages, and have the highest expression level at S44 (metamorphosis climax) (Figure 3C). The *SFTPs* play critical roles in reducing the surface tension of pulmonary alveoli and conferring innate immunity (Mikerov et al., 2008; Weaver and Conkright, 2001), while the oxidoreductases are essential in maintaining redox homeostasis of the blood-gas barrier (Clerch and Massaro, 1992). *SOX9*, *ID4*, *KLF4*, and *TITF1* are key TFs in AECs. The expression levels of *SOX9* and *ID4* decrease with development and show the highest level at stage 41 (Figure 3D). *SOX9* is an epithelial progenitor marker, with roles in regulating epithelial cell proliferation and differentiation, and lung branching morphogenesis (Danopoulos et al., 2018; Nichane et al., 2017). Unlike *SOX9* and *ID4*, the expression levels of *KLF4* and *TITF1* peak at S44 (Figure 3D). *KLF4* is a key regulator in epithelial differentiation (Tetreault et al., 2016), while *TITF1* plays a role in regulating lung development and surfactant homeostasis, and coordinating pulmonary epithelial cell differentiation (Small et al., 2000; Yuan et al., 2000). These results indicate metamorphosis climax as a critical stage for cell growth, differentiation and functional switching.

**Figure 3.**
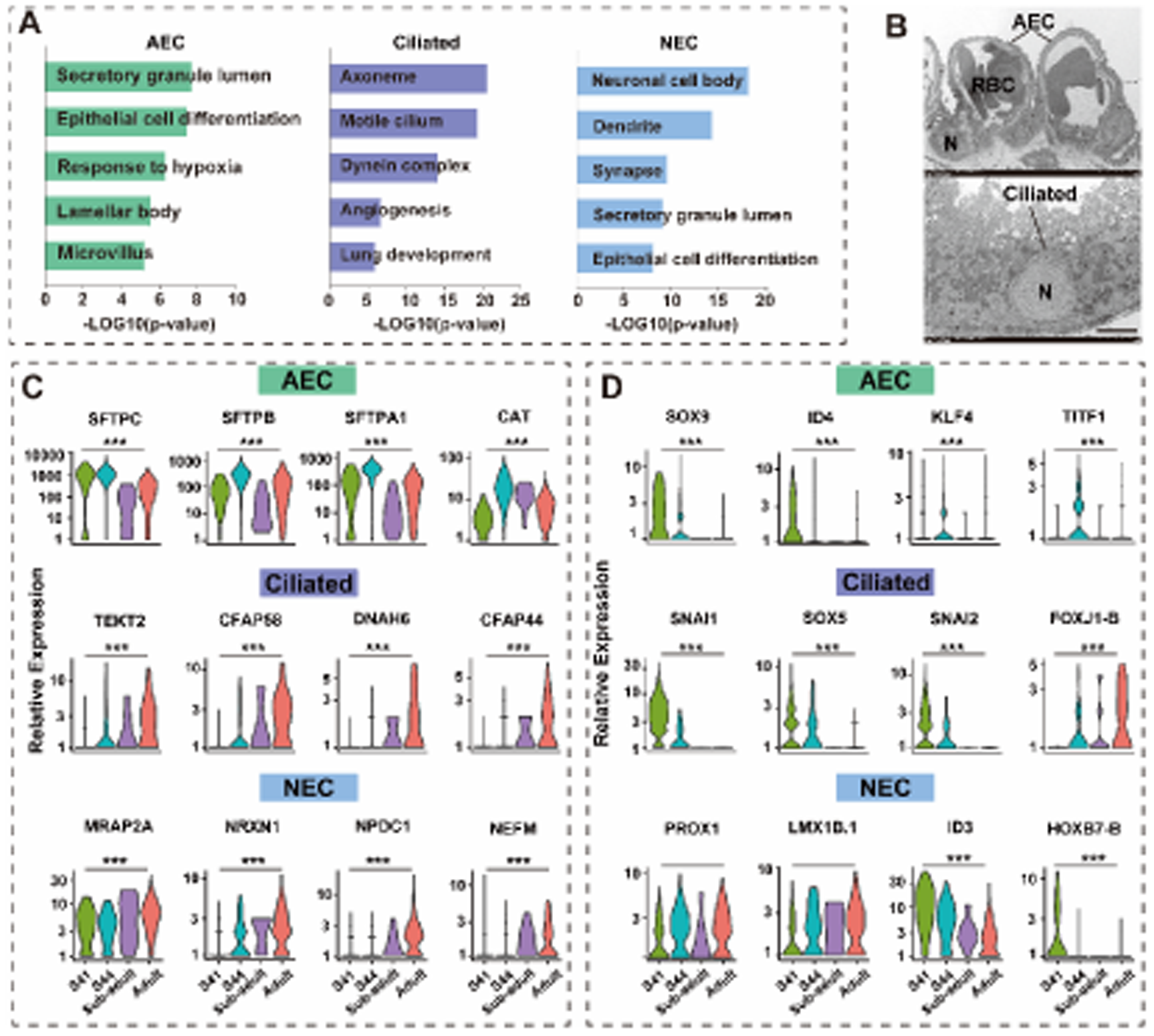
Heterogeneity and temporal molecular features of pulmonary epithelial cells in 4 developmental stages. (A) GO enrichment of feature genes in pulmonary epithelial cells (adjusted *p value* < 0.001). (B) TEM result of epithelial cell showing ultrastructural characters of AECs and ciliated cells, LB, lamellar body; MV, microvillus. Scale bar, 5 *μ*m. (C-D) Temporal dynamic expression of feature genes (C) and TFs (D) in pulmonary epithelial cells with development, (***) indicates a significant difference among the expression level in different stages (*p value* < 0.001).

Ciliated cells distributed in the tracheal epithelium were columnar and ciliated in the free surface (Figure 3B). These cells are featured by genes invovled in ciliary assembly and lung development (Figure 3A). The expression level of ciliary genes (i.e., *TEKT2*, *CFAP58*, *DNAH6*, and *CFAP44*) increases with the maturation of lung (Figure 3D). Correspondingly, *FOXJ1-B*, the key TF for motile ciliogenesis and formation of motile cilia, exhibit similar temporal expression pattern (Didon et al., 2013; Pohl and Knöchel, 2005). In contrast, the expression level of TFs regulating cell growth or differentiation (i.e., *SNAI1*, *SNAI2*, and *SOX5*) decreases with the development (Figure 3D). These results suggest a shift in the major mission of ciliated cells from self-proliferation to functional maturation during water-to-land transition.

The marker genes of NECs were enriched in neuronal differentiation, cell secretion, and epithelial cell differentiation (Figure 3A). The expression level of neuroendocrine-related genes (e.g., *MRAP2A*, *NRXN1*, *NPDC1*, and *NEFM*) increases with development (Figure 3C), while the expression level of *ID3*, a TF regulating cell cycle and survival of neural crest progenitors (Nichane et al., 2008), decreases with development (Figure 3D). The expression level of *PROX1* and *LMX1B.1*, the key regulator of neurogenesis (Demarque and Spitzer, 2010; Stergiopoulos et al., 2015) is comparable between the four developmental stages. It seems that the lungs of amphibians maintain the ability of neurogenesis even in adult stages.

These results indicate that epithelial cells keep robust cell proliferating and differentiating at S41, in favor of the rapid lung morphogenesis. Subsequently, AECs boosts the expression of pulmonary surfactant proteins at S44, a vital transitional stage from water to land. This may be a critical cellular event for the initial of air breathing. Then, the maturation of ciliated cells and NECs in juvenile and adult stages enables more advanced function of air breathing.

### Construction of differentiation trajectories of epithelial cells

AEC progenitors were identified in *M. fissipes* lungs supported by the co-expression of the AEC markers (*SFTPB*) and pulmonary stem cell marker *SOX9* (Supplementary Figure 3). Compared with AECs, AEC progenitor additionally expressed TFs for cell growth and differentiation (Figure 4A). RNA fluorescence in situ hybridization (FISH) indicates that AEC progenitors are scattered among AECs and they are identified in all the four developmental stages (Figure 4B). Differentiation trajectory analysis of AECs shows that AEC progenitor and AECs are distributed in the root and tip of the differentiation trajectory, respectively (Figure 4C). In detail, the expression of genes involving in cell proliferation and differentiation (*CCND1*, *PCNA*, *HES4*, *EGFR*, *HMCN1*, *SOX9*, *EGR1*, *MCM5A*) decreases with pseudotime, while the expression of mature AECs marker (*SFTPA1*, *SFTPB*, *SFTPC*, and *NPC*) increases with it (Supplementary Figure 3D & E). Notably, we identified two cell states of AECs (Figure 4C). State 1 cells appear since S41 and thrive at S44, and then almost disappear in sub-adult and adult lungs; state 2 cells exist in all the developmental stages (Figure 4C). Interestingly, the AEC state 1 cells were highly expressed the genes that involved in cell growth, proliferation, and differentiation potencies, such as *SFRP2*, *WNT4*, *WNT10B*, and *WNT11* with the responsible for the activation of WNT signal pathway. In contrast, the state 2 cells express genes responsible for the mature functions of AECs (Figure 4F).

**Figure 4.**
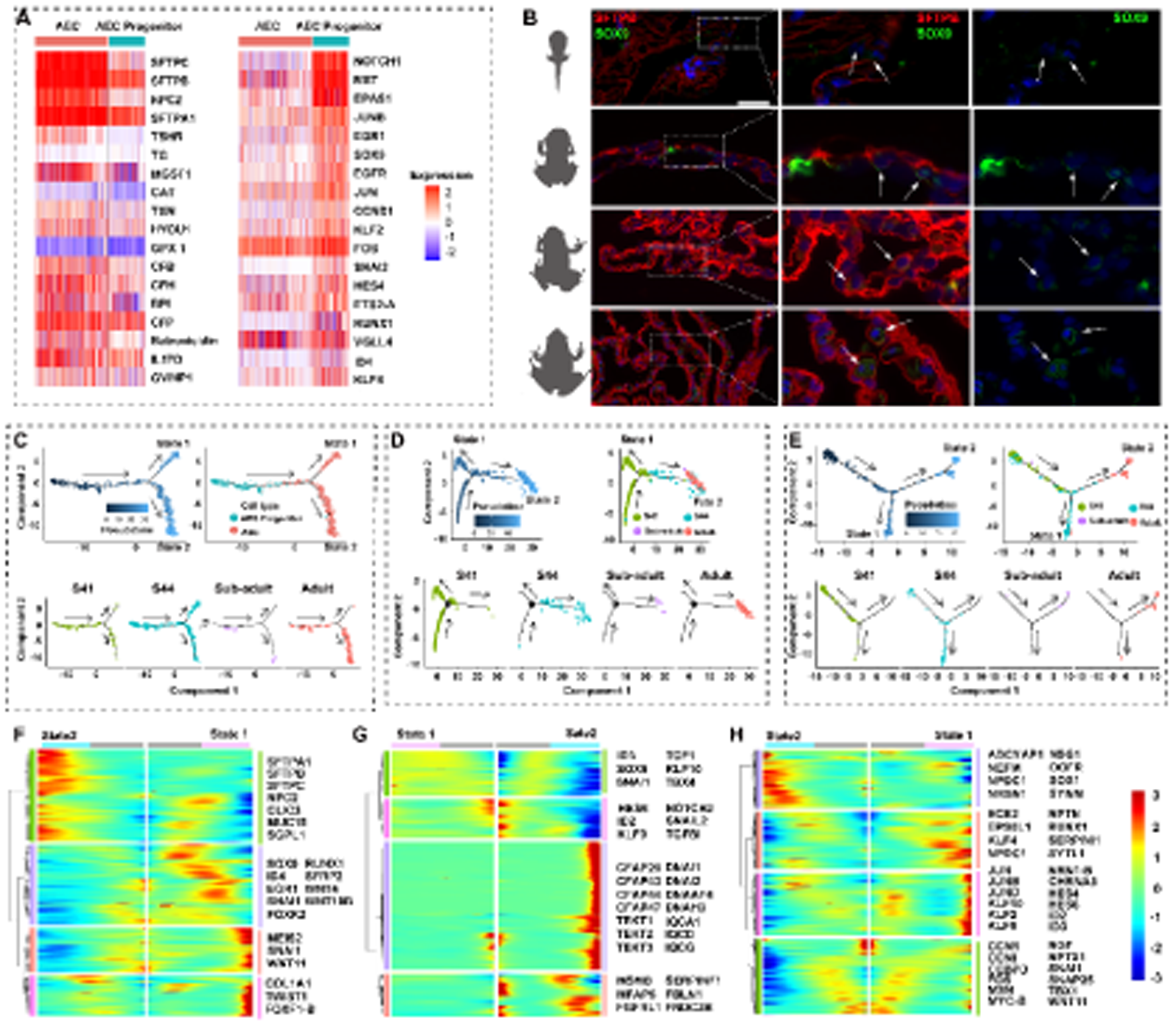
The differentiation trajectories of epithelial cells during lung development. (A) Differential expression of genes and TFs between AECs and AEC progenitors. (B) Fluorescence in situ hybridization (FISH) showing the transcriptional location of *SFTPB* (red) and *SOX9* (green) in alveoli. Scale bar, 50 *μ*m. (C-E) Pseudotime differentiation trajectory of AECs (C), Ciliated cells(D), and NECs (E). (F-H) Gene expression patterns of different cell differentiation states in AECs (F), Ciliated cells (G), and NECs (H).

Regarding to the ciliated cells or NECs, we also found two cell states according to the differentiation trajectory analysis (Figure 4D, E; Supplementary Figure 4). For ciliated cells, the state 1 cells are only identified in the lung of S41 individuals, while state 2 cells exist consistently across the four developmental stages (Figure 4D). For NECs, the state 1 cells appear since S41 and thrive at S44, and then almost disappear in sub-adult and adult lungs, the same variation pattern to that of ACE state 1. The NECs state 2 cells are identified in S44, sub-adult, and adult individuals (Figure 4E). Similar to ACEs, genes and TFs involving in cell growth, proliferation and differentiation are intensively expressed in state 1 ciliated cells and NECs, while genes associated with specialized cellular function are highly expressed in state 2 ciliated cells and NECs (Figures 4G and H).

These results indicate that pulmonary epithelial cells may differentiate into a developmental-specific transitional state, which is featured by prominent proliferation and differentiation capacity. We speculate this transitional state may be a critical mechanism for the rapid morphogenesis of amphibian pulmonary alveoli during the transition from water to land.

### The peculiarity of the AECs in *M. fissipes*

We examined and compared the ultrastructure of alveoli between *M. fissipes* and *M. musculus*. In *M. musculus* alveoli, there are 2 types of AECs containing the squamous AEC1 and the small-sized AEC2 attaching to the surface of AEC1. The AEC2 of *M. musculus* are filled with lamellar bodies in their cytoplasm and covered by microvilli on their surface (Figure 5A). However, the alveoli of *M. fissipes* were composed of only one type of squamous AECs, which is filled with lamellar in the cytoplasm and covered with dense microvilli on the cell surface (Figure 5B). These results suggest that the AECs of *M. fissipes* have the combined structural characteristics of *M. musculus* AEC1 and AEC2. Furthermore, the *M. fissipes* AECs simultaneously expressed the markers of *M. musculus* AEC1 (*AHNAK*, *MSN*, *COV*, and *LGAS3*) and AEC2 (*SFTPB*) based on the FISH and scRNA-seq data (Figure 5C & D).

**Figure 5.**
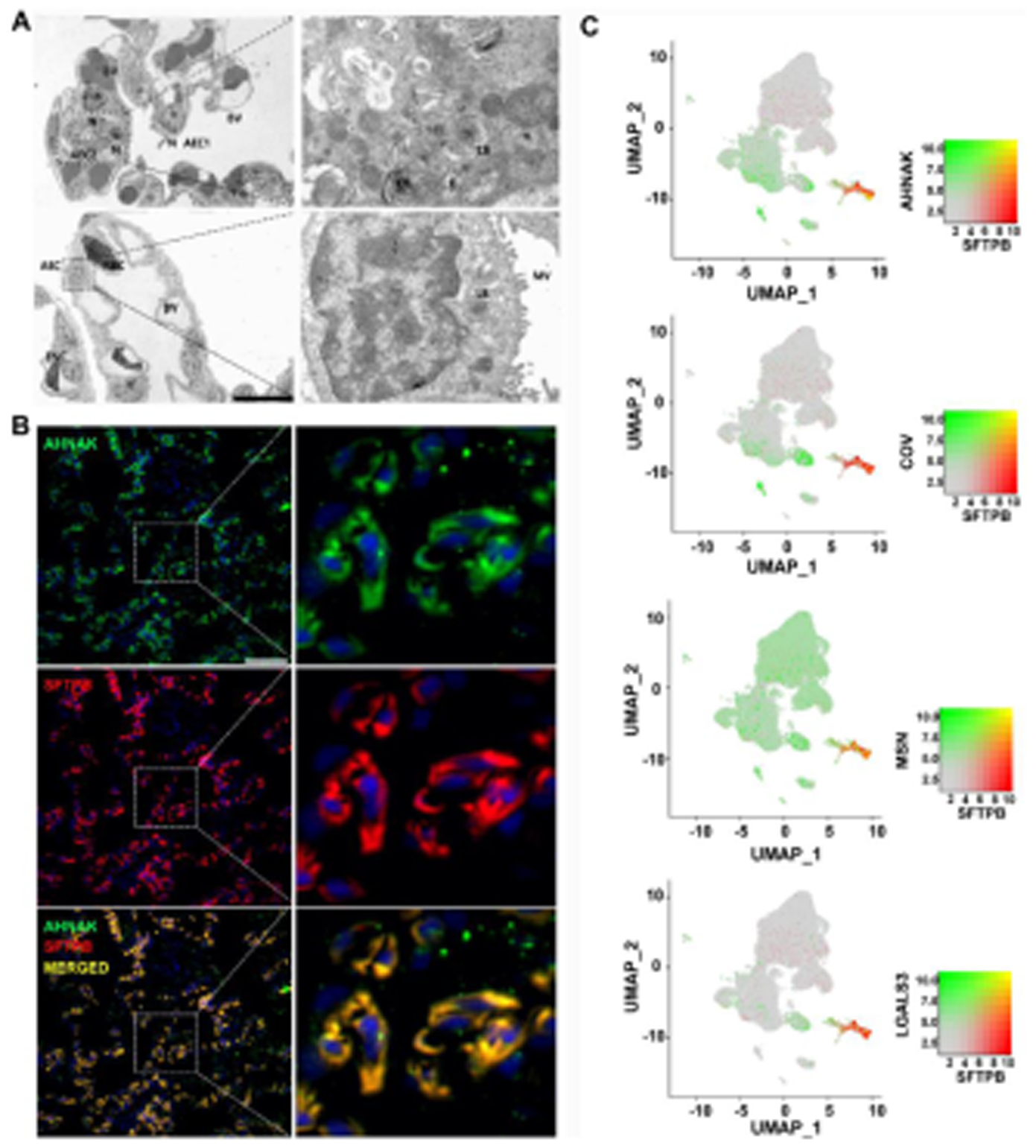
The peculiarity of AECs in adult *M. fissipes*. (A-B) Ultrastructural characters of alveoli in adult *M. fissipes* (A) and *M. musculus* (B). BV, blood vessel; LB, lamellar body; MV, microvillus; N, nucleus, scale bar, 10 *μ*m. (C) FISH presenting the expression of *AHNAK* (green) and *SFTPB* (red) in AECs of *M. fissipes*, co-expression showed yellow fluorescence. Scale bar, 50 *μ*m. (D) The UMAP plots presenting co-expression of *AHNAK*, *MSN*, *COV*, and *LGAS3* (green) with *SFTPB* (red) in AECs of *M. fissipes* based on scRNA-seq data.

### Temporal molecular features and differentiation trajectory of ECs

One type of EC was identified in *M. fissipes* lungs (Supplementary Figure 5A). These cells were characterized by cellular functions including angiogenesis, platelet degranulation, response to wounding, and establishment of endothelial barrier (Supplementary Figure 5B). The temporal expression patterns of featured genes and TFs were analyzed for ECs. In detail, the genes decreasing with development are mainly those involved in EC adhesion and migration (*VCAM1*, *PECAM*, *CLEC4M*, and *ENG*), as well as TFs (*ETS2-A*, *TEK*, *ZYX*, *EGR1*, *ID3*, *JUN*, and *JUNB*) regulating cell growth; while those increasing with development are mainly involved in angiogenesis (*CLEC14A, PROX1*, *NR2F2*, *KLF4*, and *TBX1*) and thrombin (*MMRN2* and *F5*) (Figures 6A & B). Notably, *BMP3*, *WNT10B*, and *IGFBP5* show increased expression with development, suggesting multiple signaling pathways participated in regulating angiogenesis (Figure 6A).

**Figure 6.**
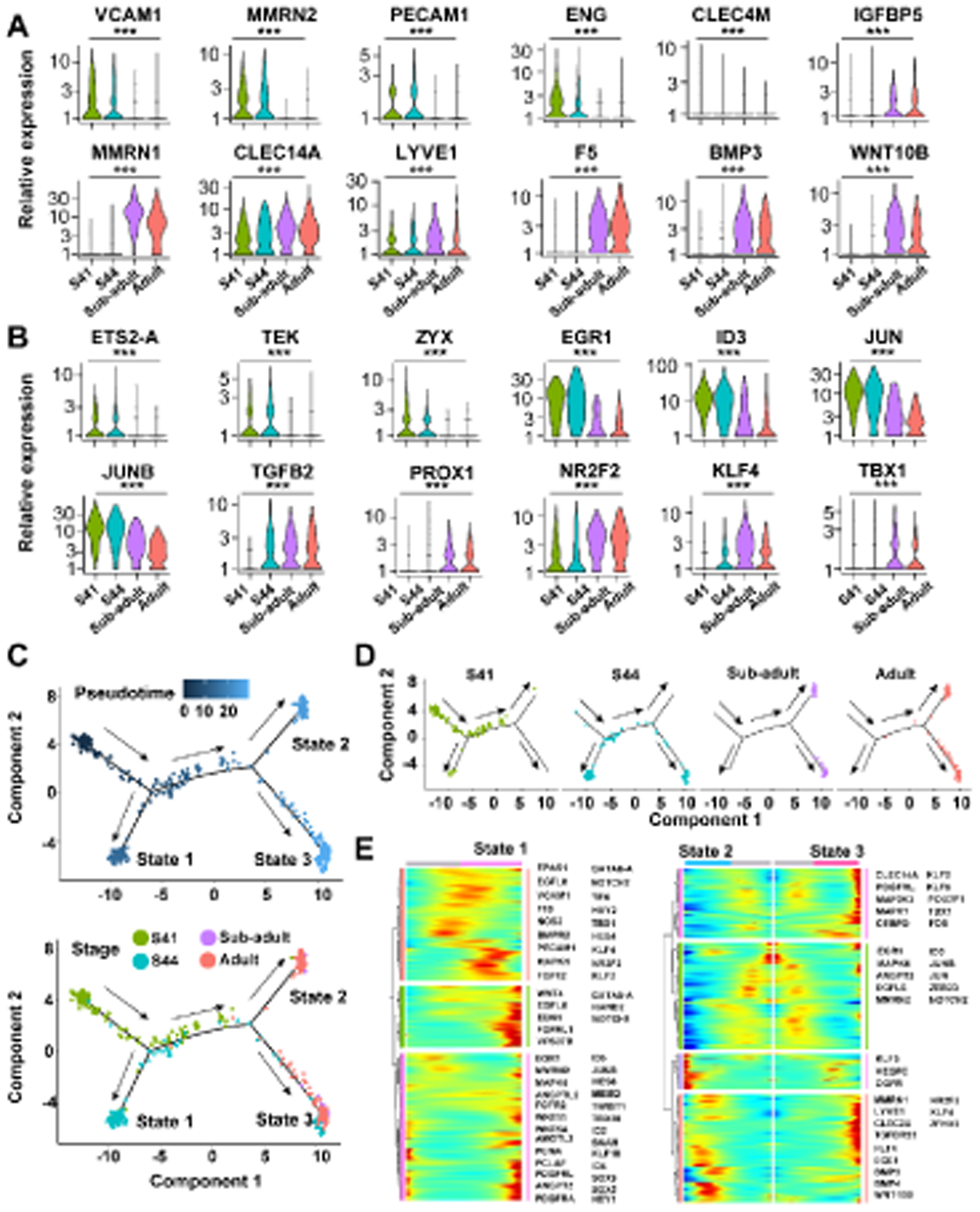
Temporal molecular features and differentiation trajectories of ECs. (A-B) Temporal dynamic expression of featured genes (A) and TFs (B) in pulmonary ECs. (***) indicates a significant difference among the expression level in different stages (p value < 0.001). (C-D) Pseudotime differentiation trajectory of ECs, dots colored by pseudotime and developmental stage (C), UMAP plots splitted by developmental stage (D). (E) Gene expression patterns of cell differentiation states in ECs.

The differentiation trajectory of ECs was constructed and 3 cell states were identified (Figures 6C). In detail, the state 1 cells increase from S41 to S44, and then disappears in sub-adult and adult lungs. State 2 cells were identified in sub-adult and adult lungs whereas state 3 cells were identified at S44, sub-adult, and adult stages (Figure 6D). The genes and TFs involving in cell growth, proliferation and differentiation show higher expression levels in state 1 cells, while those involving in angiogenesis of the lymphatic vascular systems (e.g., *VEGFC* and *KLF5*) and vascular endothelial development (e.g., *KLF2* and *KLF4*) (Bui et al., 2016; Guo *et al*., 2008; Kukk et al., 1996) show higher expression levels in state 2 and 3 cells, respectively (Figure 6E). These results suggest that state 1 cells possess considerable ability of proliferation and differentiation. It may promote rapid morphogenesis of capillary network during metamorphic climax. While state 2 and state 3 cells likely correspond to mature lymphatic and vascular ECs, respectively, and their variation of differentiation trajectory suggest that the differentiation of vascular ECs are prior to lymphatic ECs.

### Dynamics of RBCs and molecular switches of hemoglobin (HB)

Four distinct types of RBCs were identified (Figure 1A). The abundance of RBC-1, RBC-2, and RBC-4 approximately increases with development whereas that of RBC-3 decreases with it (Figure 7A). Interestingly, RBC-1 and RBC-2 exist acorss all the four stages, RBC-3 only exists in larval stages and disappears in sub-adult and adult, while RBC-4 first appeared at S44 and peaked in the adult stage (Figure 7A). These results suggest that RBC-3 and RBC-4 are likely the larval- and adult-specific cell type, respectively. The numbers of RBCs circulating through the alveoli increases with development, and this is supported by the ultrastructural dynamics of the alveoli (Figure 7B). This suggests that the pulmonary blood flow increase with the development and maturation of frog lung. Eight types of HB subunits are expressed in the pulmonary circulating RBCs, including four larval-specific types (*HBA5.L*, *HBA-L8.L*, *HBB2.L*, and *HBB2.L2*), three adult-specific types (*HBA5.A*, *HBA-L8.A*, and *HBB2.A)*, and one universal type (*HB-alpha-like*) (Figure 7C). The development of *M. fissipes* is companied by a remarkable transcriptional switch in these HBs. Specifically, larval types of HBs predominated at S41 or both at S41 and S44, then replaced by adult types after S44 (Figure 7C). These results indicate that the transition from water to land is not only accompanied by the switch of RBC types, but also a transcriptional switch in HBs in the same RBC types.

**Figure 7.**
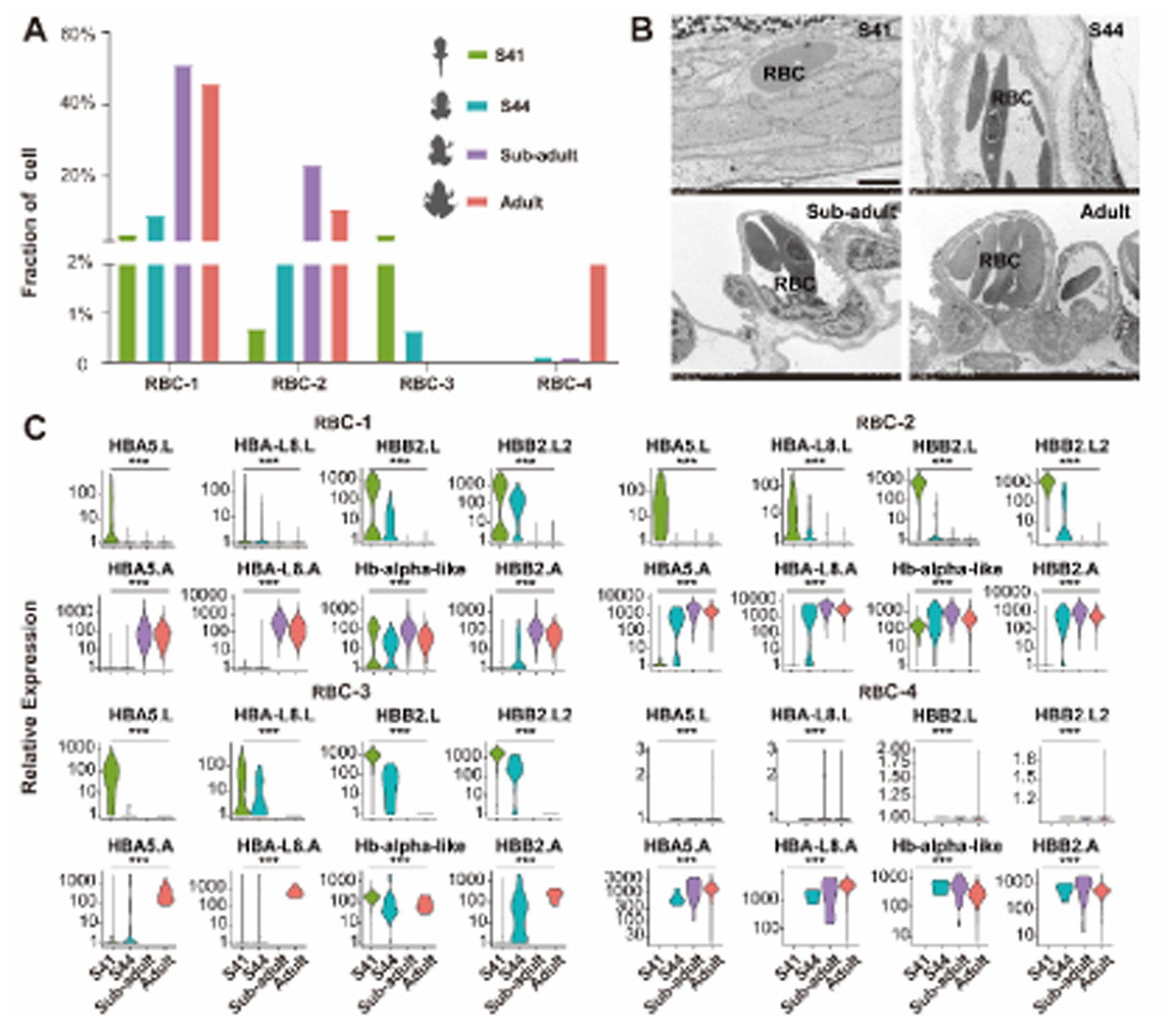
Dynamics of RBCs and molecular switches of HB in 4 developmental stages. (A) Relative proportion dynamics of RBCs. (B) Ultrastructural dynamics of the alveoli showing the proportional variations of RBCs. Scale bar, 5 *μ*m. (C) molecular switches of HB subunits in RBCs. (***) indicates a significant difference among the expression level in different stages (*p value* < 0.001).

## Discussion

In this study, we analyzed the cell heterogeneity in the lung and clarified the classification of AECs in amphibians. In addition, we elucidated the temporal dynamic features of epithelial and endothelial in developing lungs. To better understand these processes, we constructed the differentiation trajectories for epithelial and endothelial cells and revealed novel developmental-specific transition cell states. Furthermore, we illustrated the cellular and molecular processes during lung functionalization. Some thought-provoking clues were brought from the results.

The classification of AEC types in land-dwelling amphibians remains to be elucidated. We conducted a comparative analysis of AECs between *M. fissipes* and *M. musculus* at multiple biological levels and proved the existences of only one type of mature AECs in *M. fissipes*. The AEC in amphibian exhibits the combined characteristics of mammal AEC1 and AEC2. More importantly, we identified a potential AEC progenitor in *M. fissipes* based on scRNA-seq analysis. This AEC progenitor in amphibian is characterized by their high expression of *SOX9* (Figures 4A & B), and it differs from AEC progenitor for that the latter exhibits a higher degree of diversity during development (Whitsett *et al*., 2019). *M. musculus* have multiple AEC progenitors with varying differentiation potentials, and they differentiate to ACE1 and ACE2 cells at the saccular and canalicular-saccular periods (Desai et al., 2014; Frank et al., 2016). As postnatal alveolarization proceeds, mature ACE2 cells serve as the primary epithelial progenitors, and they may transdifferentiate to ACE1 during the alveolar damage repairment (Frank *et al*., 2016). For *M. fissipes*, the AEC progenitors persistently exists in the alveoli from the larval to adult stages (Figures 4B). This is suggestive of that the AEC progenitors in *M. fissipes* may have dual roles in both alveologenesis and mature alveoli regeneration after injury. These findings shed some light on the primitive state of the air-breathing organs in amphibians. The evolution of different types of AECs and AEC progenitors in amniotes may be an advanced adaptation to terrestrial life, especially when their skin became unable to assist respiration. These results also support the notion that the evolutionary divergence of the vertebrate respiratory organ is primarily driven by cell types rather than tissues (Jiang et al., 2021; Liao *et al*., 2022; Wang et al., 2021).

The temporal gene expression suggests that the epithelial and endothelial cells of *M. fissipes* lung undergo a shift in the major mission from self-proliferating and differentiating to functional maturating and specialization during water-to-land transition. Similar to previous studies (Durmaz and Scott, 2022), we observed prominent non-linearity of cell differentiation trajectories for both epithelial and endothelial cells. And we successfully identified both the transition and mature cell states for AECs, ciliated cells, NECs, and ECs (Figures 4C-E & Figures 6C-D). Interestingly, these transition cell states, which have proliferating and differentiating potentials, always boost in number at S41 and/or S44, the critical water-to-land transitional stages. Given that amphibians complete the water-to-land transition within a few days (Wang *et al*., 2017), and the organogenesis requires robust cell proliferation, we may speculate that these transition cell states constitute a strategy for the rapid morphogenesis of the blood-air barrier for air breathing during metamorphic climax. Notably, the transition state of AECs, NECs, and ECs are identified in S41 and increased in S44, while that of ciliated cells are only identified in S41 (Figures 4C-E). It is suggested the differentiation of ciliated cells from trachea are prior to AECs, NECs, and ECs from pulmonary alveoli. In addition, we identified two mature cell states in ECs: vascular ECs and lymphatic ECs. Vascular ECs appeared since S44, while lymphatic ECs were initially identified in sub-adult (Figure 6C, D). It indicates that although vascular ECs and lymphatic ECs originate from a common progenitor cell, the differentiation of vascular ECs occurs prior to that of lymphatic ECs.

Whilst the lungs of frog commence ventilation at pre-metamorphosis, a significant reconfiguration of their pulmonary tissue structure and gene expression is observed at metamorphotic climax (Burggren and Infantino, 2015; Chang *et al*., 2022). It is considered a hallmark of functionalization for air breathing, yet now the knowledge about the cellular and molecular dynamic changes in this process are limited. Here, we systematically revealed multiple cell types, from the conducting airways to peripheral saccules and alveoli, are involved in the functionalization of frog lung during the metamorphosis. MCs are the predominant major cell type during the tadpole stage, and diverse MCs provides the architectural niche regions necessary for lung morphogenesis and functionalization (Figures 2). As metamorphosis progresses, the overall proportion of MCs decreases, leading to a thinner alveolar septal wall (Figures 1D, E). This process is similar to the changes of *M. muscculus* lung at birth (Guo *et al*., 2019) and the change is likely a precondition for efficient gas exchange. Epithelial cells (including AECs, ciliated cells, and NECs) and ECs maintain their ability to growth, proliferate and differentiate during the tadpole stages. In AECs, the activation of *SOX9*, *KLF4*, and *TITF1* orchestrates intricate cellular processes, encompassing growth and differentiation mechanisms (Figures 3D). Likewise, the activation of *SNAI1*, *SNAI2*, and *SOX5* in ciliated cells play indispensable roles in governing their proliferation and differentiation processes. NECs display the activation of the *ID3*, contributing significantly to their specific differentiation (Figures 3D). Furthermore, the activation of *TEK*, *ZYX*, and *EGR1* in ECs regulate the cell growth and proliferation (Figures 5B). By harmonizing the activities of these cell-specific TFs, epithelial cells and ECs intricately coordinate their growth and differentiation processes, ensuring the precise formation of the respiratory epithelium and pulmonary vascular networks. In addition, adapting for air breathing requires the activation of a series of adaptive responses to the sudden transition from the relatively hypoxic intrauterine environment to the relatively hyperoxic extrauterine environment in mammals (Burri, 1984; Cao and Kaufman, 2014). Similarly, surfactant proteins and oxidoreductases showed the high expression in AECs at metamorphosis climax, playing critical functions in reducing the surface tension of pulmonary alveoli and in maintaining redox homeostasis of the blood-gas barrier, respectively. At the subadult and adult stages, ciliary components, neuroendocrine functions, and angiogenesis related genes showed increasing expression in ciliated cells, NECs, and ECs, respectively (Figures 3 & Figures 6). It demonstrates mature cell functions, indicating the functionalization of the blood-air barrier. HBs transition from larval type to adult type in amphibian development is a physiologically important process for air breathing (Chang *et al*., 2022; Liao *et al*., 2022; Mukhi et al., 2010). Our findings indicate frog metamorphosis is accompanied not only by a switch in types of RBCs, but also by a transcriptional switch in HBs in the same types of RBC (Figures 7). These switches may be advantageous for oxygen binding and transport.

Together, these findings demonstrate a complex process by which multiple cell types work together to facilitate the functionalization of lung for air breathing.

### Limitations of the study

For scRNA-seq analysis, we isolated single cells from lungs across S41 to Adult with only two biological replicates each stage, while there is no significant difference in cell fraction between the two biological replicates from the same group/stage. It ensured the robustness of scRNA-seq data. The conclusions inferred by bioinformatics methods, such as trajectory analysis and transition cell states, require further validation by experimental and functional investigations.

## Methods

### Animals

Four egg clutches (ranging from 200 to 500 eggs) of *M. fissipes* were obtained in lab and placed into 12 aquatic containers (length 42 × width 30 × depth 10 cm, water depth = 5 cm) and hatched (water temperature 25 ± 0.5 °C, light/dark = 12:12 h, lights on at 7:00 h, off at 19:00 h). The hatched tadpoles were fed with the solution of boiled chicken egg yolk once a day for 2 days. Tadpoles were next fed with spirulina powder (China National Salt Industry Corporation) once a day, and water was replaced every 2 days. The developmental stages of tadpoles were identified according to the staging table reported by Wang et al. (2017). The adults *M. fissipes* frogs used in this work were the parents of the tadpoles and subadults above, which were collected from farmlands (E 103.459885°, N 30.744614°, 701 m) located in Shifang City, Sichuan Province, China.

All procedures applied for this study were approved by the Institutional Ethics Committee of Animal Ethical and Welfare Committee of Chengdu Institute of Biology, Chinese Academy of Sciences (permit: CIB20190201), and all methods were carried out in accordance with the Code of Practice for the Care and Handling of animal guidelines. The study is reported in compliance with the ARRIVE guidelines.

### Tissue dissociation and preparation of single-cell suspensions

For scRNA-seq, 90, 70, 30, and 5 individual lungs at stage 41, stage 44, subadult (2 months after hatching), and adult, respectively were dissociated for preparation of single-cell suspensions. Two biological replicates were prepared for each developmental stage. The lung tissues were placed into the culture dish on ice with 0.7 × HBSS (calcium-free and magnesium-free), and cut them into 0.5 mm2 pieces, followed by 0.7 × HBSS washed. Lung tissues were grinded with ground glasses and were transferred in dissociation solution (0.35% collagenase IV5, 2 mg/ml papain, 120 Units/ml DNase I) on ice for 15-20 min and were pipetted 4-5 times with a Pasteur pipette. The tissues were washed with 0.7 × HBSS, and dissociated with trypsin in 34 ℃ water bath for 8 min. Digestion was terminated with 0.7× HBSS containing 10% fetal bovine serum (FBS, V/V). The cell suspension was pipetted 5-10 times with a Pasteur pipette and filtered by passing through 30 *μ*m stacked cell strainer and centrifuged at 300g for 5 min at 4 °C. The cell pellets were resuspended in 100 *μ*l 0.7× HBSS (0.04% BSA). The suspension was next centrifuged at 300g for 5 min at room temperature and resuspended in 100 *μ*l Dead Cell Removal MicroBeads (MACS 130-090-101), followed by removed the dead cells using Miltenyi ® Dead Cell Removal Kit (MACS 130-090-101). The suspension was furtherly resuspended in 1× HBSS (0.04% BSA) and centrifuged at 300 g for 3 min at 4 °C (repeat 2 times). The cell pellets were resuspended in 50 *μ*l of 0.7 × PBS (0.04% BSA). The overall cell viability was confirmed by trypan blue exclusion and the percentage of viable cells with greater than 85% were chosen for further analyzed. Single cell pellets were counted using a Countess II Automated Cell Counter and concentration adjusted to 700-1200 cells/*μ*l.

### Chromium 10× Genomics library and sequencing

scRNA-seq libraries were prepared following the manufacturer’s instructions of Chromium Next GEM Single Cell 3ʹ Reagent Kits v3.1 (https://www.10xgenomics.com/support/single-cell-gene-expression/documentation/steps/library-prep/chromium-single-cell-3-reagent-kits-user-guide-v-3-1-chemistry Chemistry) from 10× Genomics, Inc. (Pleasanton, CA). Libraries were sequenced on Illumina NovaSeq 6000 sequencing system (paired-end multiplexing run,150bp) by LC-Bio Technology Co. Ltd. (Hangzhou, China).

### Pre-processing and quality control of scRNA-seq

Sequencing results were demultiplexed and converted to FASTQ format using Illumina bcl2fastq software (version 2.20). Sample demultiplexing, barcode processing and single-cell 3’gene counting using the Cell Ranger pipeline (https://support.10xgenomics.com/single-cell-geneexpression/software/pipelines/latest/what-is-cell-ranger, version 3.1.0) and scRNA-seq data were aligned to *M. fissipes* reference genome (unpublished). The Cell Ranger output was loaded into Seurat (version 4.0.2). High quality single cells must pass the quality control threshold: the number of detected genes per cell > 200 and < 3000 (all genes expressed in less than three cells were removed), the percent of mitochondrial-DNA derived gene-expression < 5%.

### Integration of scRNAseq datasets from four developmental stages

Datasets from eight sequencing libraries were integrated using anchoring procedure implemented in Seurat (version 4.0.2). In details, data was normalized with the fast integration using reciprocal PCA (RPCA) method using the “NormalizeData” function, and subsequently scaled by the Pearson Residuals with a scale factor of 10,000. The top 3000 highly variable features were selected using the “SelectIntegrationFeatures” function, followed by finding the integration anchors using the “FindIntegrationAnchors” function, performing the integration of the data using the “IntegrateData” function.

### Identification of cell clusters

Following integration, principal component analysis was performed using the “RunPCA” function with default parameters, UMAP dimensionality reduction methods were conducted based on the top 50 principal components (PCs) using the “RunTSNE” and “RunUMAP” functions, respectively. Moreover, unsupervised clusters were identified by setting the top 50 PCs and a clustering resolution of 0.35 using “FindNeighbors” and “FindClusters” functions.

### Identification of differentially expressed genes (DEGs) across clusters

FindAllMarkers function implemented in Seurat (version 4.0.2) was used to identify DEGs across clusters with the options “min.pct = 0.25, logfc.threshold = 0.25, test.use = wilcox”. Multiple test correction for *p* value was performed using the Bonferroni method, and 0.05 was set as a threshold to define significance.

### Gene ontology (GO) enrichment analysis

Gene Ontology (GO) analysis was conducted on the KOBAS 3.0. Benjamini– Hochberg (BH) method was used for the multiple test adjustment, and 0.05 was set as a threshold to define significance.

### Annotation of cell cluster

Cell clusters were annotated based on the expression levels of conservative cell markers from Cell Marker 2.0 Database and differential expression gene (DEGs) profile of cell clusters. Cell markers were visualized using “FeaturePlot” function.

### Pseudotime trajectory analysis

Monocle 2 package was used to constructed the differentiation trajectory of epithelial cell (including AECs, NECs, and ciliated cells) and ECs. Genes expressed in less than 10 cells were filtered out. DEGs were computed by function “differentialGeneTest” in monocle 2. Genes with q-value less than 0.01 were regarded as DEGs and sorted by q-value using “setOrderingFilter” function. The pseudotime trajectory was constructed by “DDRTree” algorithm with default parameters. The dynamical expression changes of selected marker genes by pseudotime were visualized by “plot_genes_in_pseudotime” and “plot_pseudotime_heatmap” function. Branches in the trajectories were analyzed and visualized by “plot_genes_branched_heatmap”.

### Transmission electron microscopic observation

The lung tissue samples were obtained from S41, S44, subadult, and adult individuals, with three biological replicates for each stage. Fresh lung tissues were collected and cut into small blocks (1 mm³). These tissue blocks were fixed in 3% glutaraldehyde at 4°C for 6 hours, followed by rinsing in 0.1 M Sorensen’s phosphate buffer (pH 7.4) three times. Subsequently, the blocks were postfixed in 1% osmium tetroxide in the same buffer for 2 hours. To proceed, the tissue blocks were dehydrated using a graded series of ethanol (30%, 50%, 70%, 80%, 95%, and 100%) for 20 minutes each concentration. Afterward, the blocks were penetrated with a mixture of acetone and EMBed 812 overnight at 37°C. The tissue blocks were then embedded in EMBed 812, and the embedding models with resin and samples were polymerized at 65°C for more than 48 hours. Next, the resin blocks were cut into ultrathin sections measuring 60–80 nm using an ultramicrotome (Leica UC7) and Diamond slicer (Daitome Ultra 45). These ultrathin sections were placed onto 150-mesh cuprum grids with formvar film and stained in a solution of 2% uranium acetate saturated alcohol solution for 8 min. Following rinsing with 70% ethanol and ultrapure water, the ultrathin sections were stained with 2.6% lead citrate for 8 minutes. After drying with filter paper, the cuprum grids were placed into a grid board overnight at room temperature. Finally, the ultrastructure of lung was observed under a TEM (Hitachi, HT7800/HT7700) operating at 60 kV, and images were captured using a CCD digital camera (APTINA CMOS Sensor, San Jose, CA, USA).

### RNA fluorescence in situ hybridization (FISH) assay

The lung tissue samples obtained from S41, S44, subadult, and adult for RNA FISH (three biological replicates each stage). Fluorescence-conjugated probes of SFTPB, AHNAK, SOX9 for FISH were generated in line with the protocols of Servicebio Technologies and probe sequences were provided in Supplementary Data 3. A detailed step-by-step RNA FISH protocol used in this paper can be found in the manuals of Servicebio Technologies.

### Statistics and reproducibility

Gene different expression analysis across clusters using Wilcox test. Statistical significance was performed with multiple test correction for *p value* (Bonferroni). Benjamini–Hochberg (BH) method was used for the multiple test adjustment for the gene ontology enrichment analysis. Gene different expression analysis across clusters using Kruskal-Wallis test. Unless stated elsewhere, all FISH and TEM were performed with at least three experimental replicates. No statistical method was used to predetermine sample size. No data were excluded from the analyses. Filtering criteria for the low-quality cells are provided in the method above. Randomization is not related to this study. The investigators were not blinded to allocation during the experiments and the outcome assessment.

## Data availability

All data needed to evaluate the conclusions in the paper are available in the GSA (CRA010691) or present in the paper and/or the Supplementary Materials. Additional data related to this paper may be requested from the authors.

## Author Contributions

Conceptualization, J.P.J., W.Z., and L.M.C.; methodology, J.P.J., W.Z., and L.M.C.; software, L.M.C. and W.Z.; validation, L.M.C. and W.Z.; formal analysis, L.M.C. and W.Z.; investigation, L.M.C., W.Z., Q.H.C., and B.W.; resources, L.M.C., W.Z., Q.H.C., M.H.Z., B.W., and J.Y.L.; data curation, L.M.C. and W.Z.; writing—original draft preparation, L.M.C.; writing—review and editing, L.M.C., W.Z. and J.P.J.; visualization, L.M.C. and W.Z.; supervision, J.P.J.; project administration, J.P.J.; funding acquisition, J.P.J. and W.Z. All authors participated in the manuscript preparation and approved the final version of the manuscript.

---

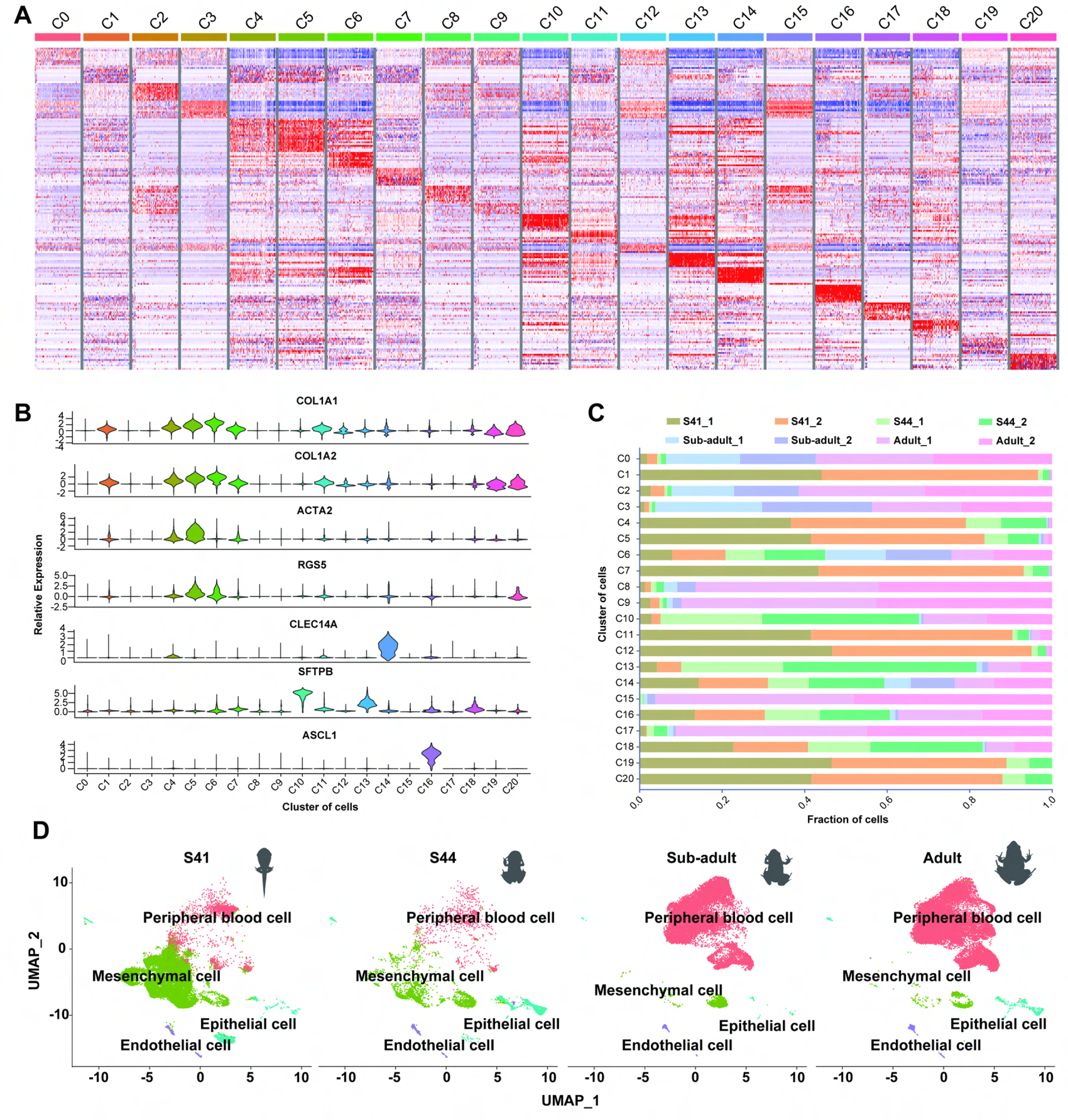

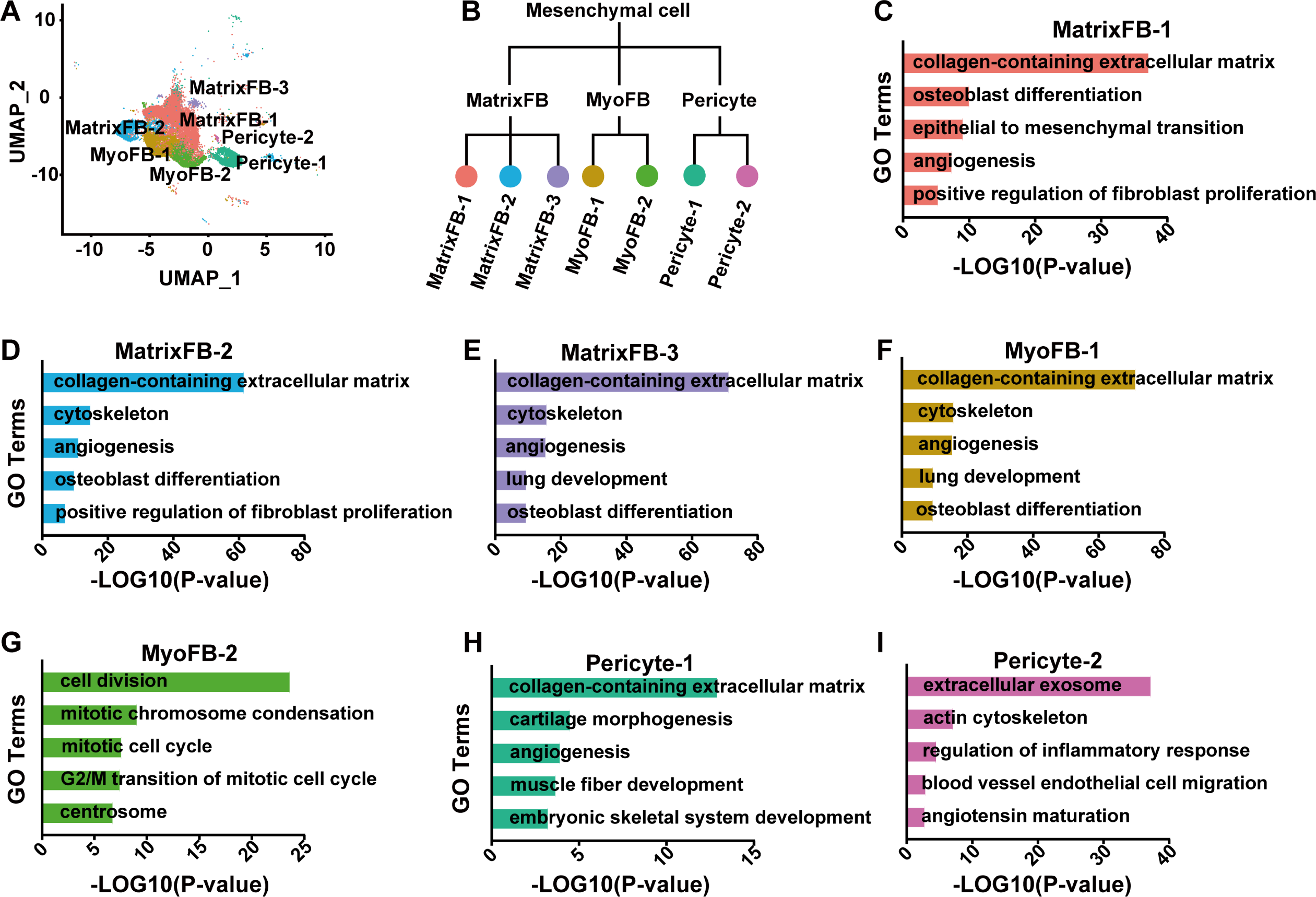

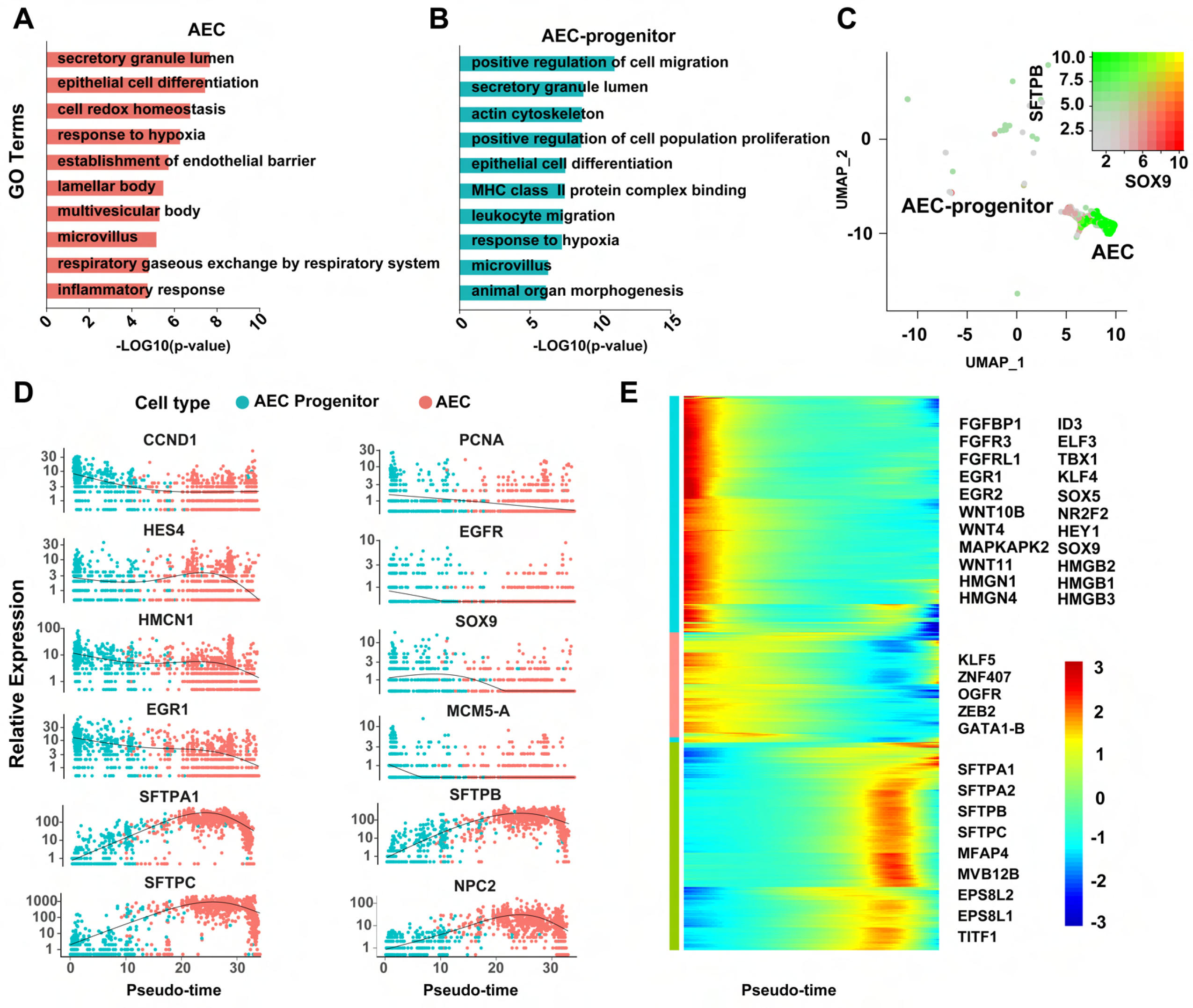

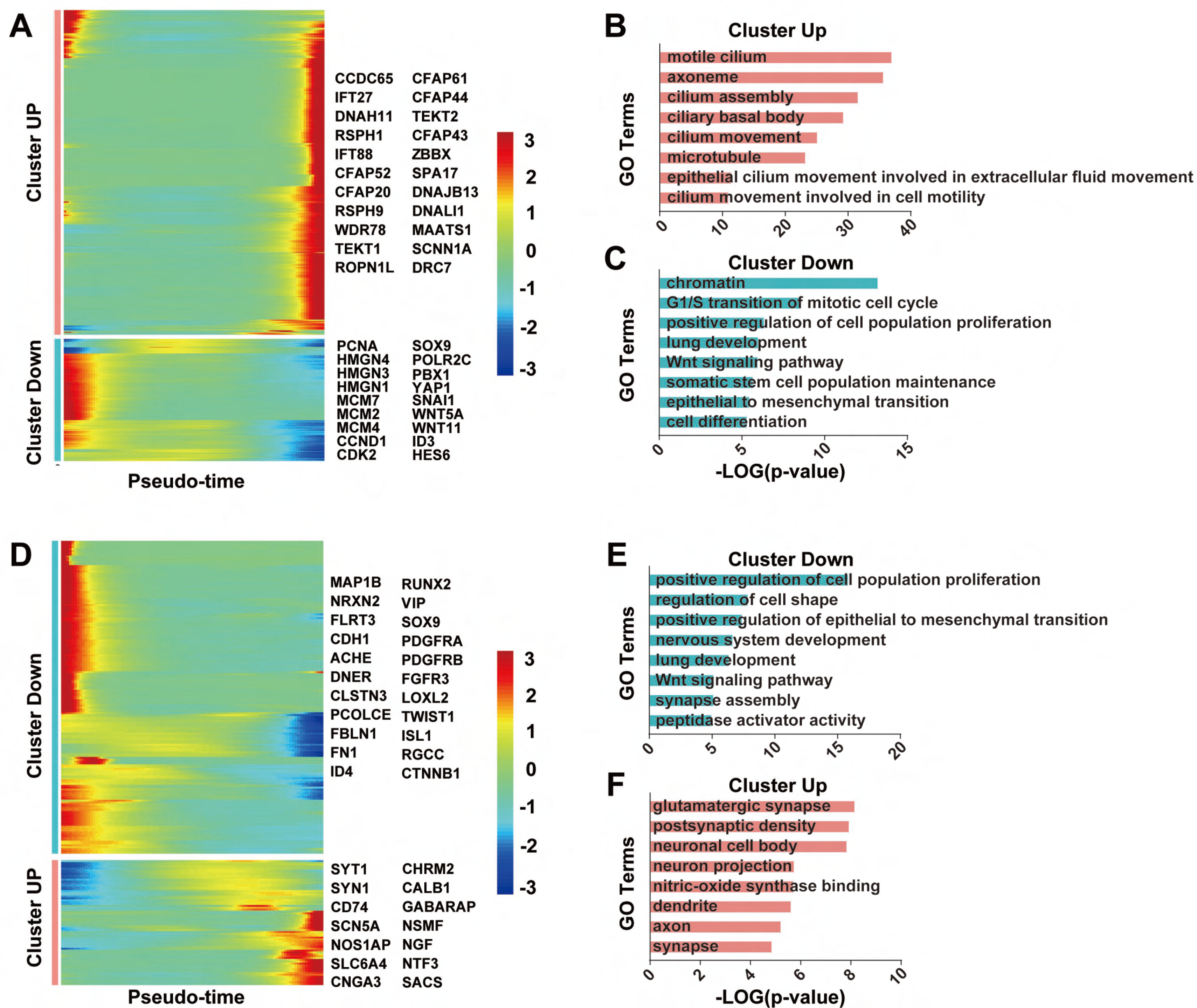

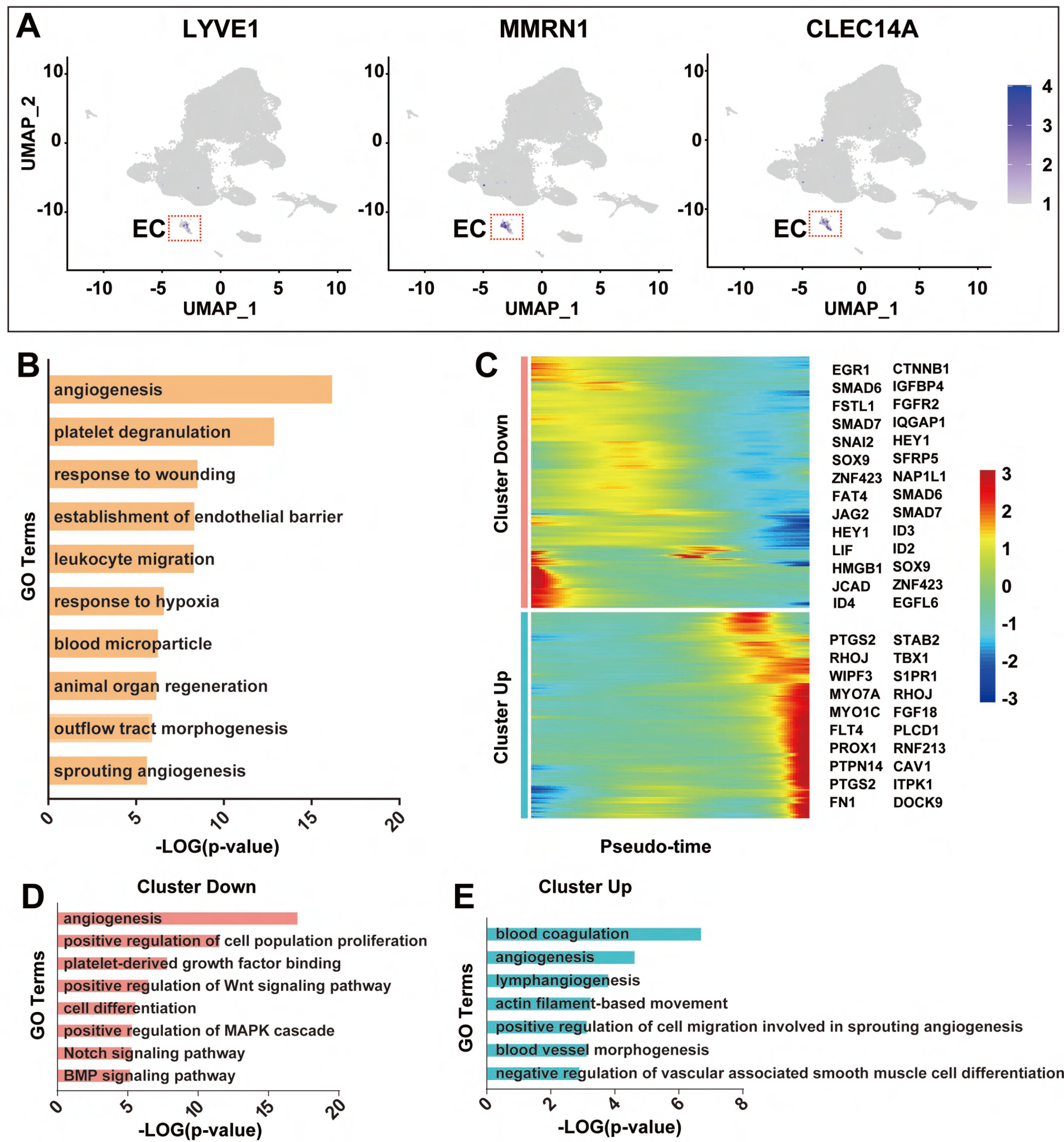

## References

Abraham, A.B., Bronstein, R., Reddy, A.S., Maletic-Savatic, M., Aguirre, A., and Tsirka, S.E. (2013). Aberrant neural stem cell proliferation and increased adult neurogenesis in mice lacking chromatin protein HMGB2. Plos One 8, e84838.

Adams, T.S., Schupp, J.C., Poli, S., Ayaub, E.A., Neumark, N., Ahangari, F., Chu, S.G., Raby, B.A., DeIuliis, G., Januszyk, M., et al. (2020). Single-cell RNA-seq reveals ectopic and aberrant lung-resident cell populations in idiopathic pulmonary fibrosis. Science Advances 6, eaba1983.

Birbrair, A., Zhang, T., Wang, Z.-M., Messi, Maria L., Mintz, A., and Delbono, O. (2014). Pericytes at the intersection between tissue regeneration and pathology. Clinical Science 128, 81–93.

Bischoff, P., Trinks, A., Obermayer, B., Pett, J.P., Wiederspahn, J., Uhlitz, F., Liang, X., Lehmann, A., Jurmeister, P., Elsner, A., et al. (2021). Single-cell RNA sequencing reveals distinct tumor microenvironmental patterns in lung adenocarcinoma. Oncogene 40, 6748–6758.

Blenkinsopp, W.K. (1967). Proliferation of respiratory tract epithelium in the rat. Experimental Cell Research 46, 144–154.

Bui, H.M., Enis, D., Robciuc, M.R., Nurmi, H.J., Cohen, J., Chen, M., Yang, Y., Dhillon, V., Johnson, K., Zhang, H., et al. (2016). Proteolytic activation defines distinct lymphangiogenic mechanisms for VEGFC and VEGFD. Journal of Clinical Investigation 126, 2167–2180.

Burggren, W.W., and Infantino, R.L., JR. (2015). The respiratory transition from water to air breathing during amphibian metamorphosis. American Zoologist 34, 238–246.

Burri, P.H. (1984). Fetal and postnatal development of the lung. Annual Review of Physiology 46, 617–628.

Cao, S.S., and Kaufman, R.J. (2014). Endoplasmic reticulum stress and oxidative stress in cell fate decision and human disease. Antioxidants & Redox Signaling 21, 396–413. 10.1089/ars.2014.5851.

Chang, L.M., Zhang, M.H., Chen, Q.H., Liu, J.Y., Zhu, W., and Jiang, J.P. (2022). From water to land: the structural construction and molecular switches in lungs during metamorphosis of *Microhyla fissipes*. Biology 11, 528.

Chen, D.S., Sun, J., Zhu, J.C., Ding, X.N., Lan, T.M., Wang, X.R., Wu, W.Y., Ou, Z.H., Zhu, L.N., Ding, P.W., et al. (2021). Single cell atlas for 11 non-model mammals, reptiles and birds. Nature Communications 12, 7083.

Clerch, L.B., and Massaro, D. (1992). Oxidation reduction sensitive binding of lung protein to rat catalase messenger RNA. Journal of Biological Chemistry 267, 2853–2855.

Cui, Y.L., Zheng, Y.X., Liu, X.X., Yan, L.Y., Fan, X.Y., Yong, J., Hu, Y.Q., Dong, J., Li, Q.Q., Wu, X.L., et al. (2019). Single-cell transcriptome analysis maps the developmental track of the human heart. Cell Reports 26, 1934-+.

Danopoulos, S., Alonso, I., Thornton, M.E., Grubbs, B.H., Bellusci, S., Warburton, D., and Al Alam, D. (2018). Human lung branching morphogenesis is orchestrated by the spatiotemporal distribution of ACTA2, SOX2, and SOX9. American Journal of Physiology-Lung Cellular and Molecular Physiology 314, L144–L149.

Deb, A., Davis, B.H., Guo, J., Ni, A., Huang, J., Zhang, Z., Mu, H., and Dzau, V.J. (2007). SFRP2 regulates cardiomyogenic differentiation by inhibiting a positive transcriptional autofeedback loop of Wnt3a. Stem Cells 26, 35–44.

Demarque, M., and Spitzer, N.C. (2010). Activity-dependent expression of Lmx1b regulates specification of serotonergic neurons modulating swimming behavior. Neuron 67, 321–334.

Desai, T.J., Brownfield, D.G., and Krasnow, M.A. (2014). Alveolar progenitor and stem cells in lung development, renewal and cancer. Nature 507, 190–194.

Didon, L., Zwick, R.K., Chao, I.W., Walters, M.S., Wang, R., Hackett, N.R., and Crystal, R.G. (2013). RFX3 modulation of FOXJ1 regulation of cilia genes in the human airway epithelium. Respiratory Research 14, 70.

Dore-Duffy, P. (2008). Pericytes: pluripotent cells of the blood brain barrier. Current Pharmaceutical Design 14, 1581–1593.

Durmaz, A., and Scott, J.G. (2022). Stability of scRNA-seq analysis workflows is susceptible to preprocessing and is mitigated by regularized or supervised approaches. Evolutionary Bioinformatics 18, 11769343221123050.

Dusart, P., Hallström, B.M., Renné, T., Odeberg, J., Uhlén, M., and Butler, L.M. (2019). A systems-based map of human brain cell-type enriched genes and malignancy-associated endothelial changes. Cell Reports 29, 1690–1706.e1694.

Fleetwood, J.N., and Munnell, J.F. (1996). Morphology of the airways and lung parenchyma in hatchlings of the loggerhead sea turtle, *Caretta caretta*. Journal of Morphology 227, 289–304.

Frank, D.B., Peng, T., Zepp, J.A., Snitow, M., Vincent, T.L., Penkala, I.J., Cui, Z., Herriges, M.J., Morley, M.P., Zhou, S., et al. (2016). Emergence of a wave of Wnt signaling that regulates lung alveologenesis by controlling epithelial self-renewal and differentiation. Cell Reports 17, 2312–2325.

Fritz, A., and De Robertis, E.M. (1988). *Xenopus* homeobox-containing cDNAs expressed in early development. Nucleic Acids Research 16, 1453–1469.

Gomes, F.C.A., Sousa, V.d.O., and Romão, L. (2005). Emerging roles for TGF-β1 in nervous system development. International Journal of Developmental Neuroscience 23, 413–424.

Goniakowska-Witalińska, L. (1978). Ultrastructural and morphometric study of lung of european Salamander, *Salamander salamander* L. Cell and Tissue Research 191, 343–356.

Goniakowska-Witalińska, L. (1986). Lung of the tree frog, *Hyla arborea* L, a scanning and transmission electron microscopic study. Anatomy and Embryology 174, 379–389.

Guo, M.Z., Du, Y.N., Gokey, J.J., Ray, S., Bell, S.M., Adam, M., Sudha, P., Perl, A.K., Deshmukh, H., Potter, S.S., et al. (2019). Single cell RNA analysis identifies cellular heterogeneity and adaptive responses of the lung at birth. Nature Communications 10, 37.

Guo, P., Dong, X., Zhao, K., Sun, X., Li, Q., and Dong, J. (2008). KLF5 is an essential regulator of Myc expression. Cancer Research 68, 66–66.

He, P., Lim, K., Sun, D., Pett, J.P., Jeng, Q., Polanski, K., Dong, Z., Bolt, L., Richardson, L., Mamanova, L., et al. (2022). A human fetal lung cell atlas uncovers proximal-distal gradients of differentiation and key regulators of epithelial fates. Cell 185, 4841–4860 e4825.

Hsia, C.C.W., Schmitz, A., Lambertz, M., Perry, S.F., and Maina, J.N. (2013). Evolution of air breathing: oxygen homeostasis and the transitions from water to land and sky. Compr Physiol 3, 849–915.

Hu, C., Li, T., Xu, Y., Zhang, X., Li, F., Bai, J., Chen, J., Jiang, W., Yang, K., Ou, Q., et al. (2022). CellMarker 2.0: an updated database of manually curated cell markers in human/mouse and web tools based on scRNA-seq data. Nucleic Acids Research.

Hurley, K., Ding, J., Villacorta-Martin, C., Herriges, M.J., Jacob, A., Vedaie, M., Alysandratos, K.D., Sun, Y.L., Lin, C., Werder, R.B., et al. (2020). Reconstructed single-cell fate trajectories define lineage plasticity windows during differentiation of human PSC-derived distal lung progenitors. Cell Stem Cell 26, 593–608.e598.

Jansing, N.L., McClendon, J., Henson, P.M., Tuder, R.M., Hyde, D.M., and Zemans, R.L. (2017). Unbiased quantitation of alveolar type II to alveolar type I cell transdifferentiation during repair after lung Injury in Mice. American Journal of Respiratory Cell and Molecular Biology 57, 519–526.

Jiang, M., Xiao, Y., Ma, L., Wang, J., Chen, H., Gao, C., Liao, Y., Guo, Q., Peng, J., and Han, X. (2021). Characterization of the zebrafish cell landscape at single-cell resolution. Frontiers in Cell and Developmental Biology, 2734.

Johansen, T. (2019). Selective autophagy: RNA comes from the vault to regulate p62/SQSTM1. Current Biology 29, R297–r299.

Kadur, L., Murthy, P., Sontake, V., Tata, A., Kobayashi, Y., Macadlo, L., Okuda, K., Conchola, A.S., Nakano, S., Gregory, S., et al. (2022). Human distal lung maps and lineage hierarchies reveal a bipotent progenitor. Nature 604, 111–119.

Klemm, R.D., Gatz, R.N., Westfall, J.A., and Fedde, M.R. (1979). Microanatomy of the lung parenchyma of a tegu lizard *Tupinambis nigropunctatus*. Journal of Morphology 161, 257–279.

Kukk, E., Lymboussaki, A., Taira, S., Kaipainen, A., Jeltsch, M., Joukov, V., and Alitalo, K. (1996). VEGF-C receptor binding and pattern of expression with VEGFR-3 suggests a role in lymphatic vascular development. Development 122, 3829–3837.

Liao, Y., Ma, L., Guo, Q., E, W. Fang, X., Yang, L., Ruan, F., Wang, J., Zhang, P., Sun, Z., et al. (2022). Cell landscape of larval and adult Xenopus laevis at single-cell resolution. Nature Communications 13, 4306.

Liu, L., Zhao, L., Wang, S., and Jiang, J. (2016). Research proceedings on amphibian model organisms. Zoological Research 37, 237–245.

Maina, J.N. (1989). The morphology of the lung of the black mamba *Dendroaspis polylepis* (Reptilia: Ophidia: Elapidae). A scanning and transmission electron microscopic study. J Anat 167, 31–46.

Maina, J.N., and King, A.S. (1989). The lung of the emu, Dromaius novaehollandiae: a microscopic and morphometric study. J Anat 163, 67–73.

Metzger, R.J., Klein, O.D., Martin, G.R., and Krasnow, M.A. (2008). The branching programme of mouse lung development. Nature 453, 745–750.

Mikerov, A.N., Umstead, T.M., Gan, X., Huang, W., Guo, X., Wang, G., Phelps, D.S., and Floros, J. (2008). Impact of ozone exposure on the phagocytic activity of human surfactant protein A (SP-A) and SP-A variants. American Journal of Physiology-Lung Cellular and Molecular Physiology 294, L121–130.

Morton, S.U., and Brodsky, D. (2016). Fetal physiology and the transition to extrauterine life. Clinics in Perinatology 43, 395–407.

Mukhi, S., Cai, L., and Brown, D.D. (2010). Gene switching at Xenopus laevis metamorphosis. Dev Biol 338, 117–126.

Nakahara, Y., Hashimoto, N., Sakamoto, K., Enomoto, A., Adams, T.S., Yokoi, T., Omote, N., Poli, S., Ando, A., Wakahara, K., et al. (2021). Fibroblasts positive for meflin have anti-fibrotic properties in pulmonary fibrosis. European Respiratory Journal 58.

Nemeth, M.J., Kirby, M.R., and Bodine, D.M. (2006). Hmgb3 regulates the balance between hematopoietic stem cell self-renewal and differentiation. Proceedings of the National Academy of Sciences of the United States of America 103, 13783–13788.

Nichane, M., de Crozé, N., Ren, X., Souopgui, J., Monsoro-Burq, A.H., and Bellefroid, E.J. (2008). Hairy2-Id3 interactions play an essential role in Xenopus neural crest progenitor specification. Dev Biol 322, 355–367.

Nichane, M., Javed, A., Sivakamasundari, V., Ganesan, M., Ang, L.T., Kraus, P., Lufkin, T., Loh, K.M., and Lim, B. (2017). Isolation and 3D expansion of multipotent Sox9(+) mouse lung progenitors. Nature Methods 14, 1205-+.

Okada, Y., Ishiko, S., Daido, S., Kim, J., and Ikeda, S. (1962). Comparative morphology of the lung with special reference to the alveolar epithelial cells: I. Lung of the amphibia. Acta Tuberculosea Japonica 11, 63–72.

Perry, S.F., and Sander, M. (2004). Reconstructing the evolution of the respiratory apparatus in tetrapods. Respiratory Physiology & Neurobiology 144, 125–139.

Pike, F.H. (1924). On the difficulties encountered in the evolution of air-breathing vertebrates. Science 59, 402–403.

Plasschaert, L.W., Žilionis, R., Choo-Wing, R., Savova, V., Knehr, J., Roma, G., Klein, A.M., and Jaffe, A.B. (2018). A single-cell atlas of the airway epithelium reveals the CFTR-rich pulmonary ionocyte. Nature 560, 377–381.

Pohl, B.S., and Knöchel, W. (2005). Of fox and frogs: fox (fork head/winged helix) transcription factors in *Xenopus* development. Gene 344, 21–32.

Quaggin, S.E., Schwartz, L., Cui, S., Igarashi, P., Deimling, J., Post, M., and Rossant, J. (1999). The basic-helix-loop-helix protein Pod1 is critically important for kidney and lung organogenesis. Development 126, 5771–5783.

Rankin, S.A., Thi Tran, H., Wlizla, M., Mancini, P., Shifley, E.T., Bloor, S.D., Han, L., Vleminckx, K., Wert, S.E., and Zorn, A.M. (2015). A molecular atlas of Xenopus respiratory system development. Developmental Dynamics 244, 69–85.

Rawlins, E.L., and Hogan, B.L.M. (2008). Ciliated epithelial cell lifespan in the mouse trachea and lung. American Journal of Physiology-Lung Cellular and Molecular Physiology 295, L231–L234.

Roberts, A.B., Sporn, M.B., Assoian, R.K., Smith, J.M., Roche, N.S., Wakefield, L.M., Heine, U.I., Liotta, L.A., Falanga, V., and Kehrl, J.H. (1986). Transforming growth factor type beta: rapid induction of fibrosis and angiogenesis in vivo and stimulation of collagen formation in vitro. Proceedings of the National Academy of Sciences 83, 4167–4171.

Rudders, S., Gaspar, J., Madore, R., Voland, C., Grall, F., Patel, A., Pellacani, A., Perrella, M.A., Libermann, T.A., and Oettgen, P. (2001). ESE-1 is a novel transcriptional mediator of inflammation that interacts with NF-kappa B to regulate the inducible nitric-oxide synthase gene. Journal of Biological Chemistry 276, 3302–3309.

Small, E.M., Vokes, S.A., Garriock, R.J., Li, D.L., and Krieg, P.A. (2000). Developmental expression of the Xenopus Nkx2-1 and Nkx2-4 genes. Mechanisms of Development 96, 259–262.

Stergiopoulos, A., Elkouris, M., and Politis, P.K. (2015). Prospero-related homeobox 1 (Prox1) at the crossroads of diverse pathways during adult neural fate specification. Frontiers in Cellular Neuroscience 8, 454.

Tetreault, M.P., Weinblatt, D., Shaverdashvili, K., Yang, Y.Z., and Katz, J.P. (2016). KLF4 transcriptionally activates non-canonical WNT5A to control epithelial stratification. Scientific Reports 6, 26130.

Travaglini, K.J., Nabhan, A.N., Penland, L., Sinha, R., Gillich, A., Sit, R.V., Chang, S., Conley, S.D., Mori, Y., Seita, J., et al. (2020). A molecular cell atlas of the human lung from single-cell RNA sequencing. Nature 587, 619–625.

Wang, J., Sun, H., Jiang, M., Li, J., Zhang, P., Chen, H., Mei, Y., Fei, L., Lai, S., Han, X., et al. (2021). Tracing cell-type evolution by cross-species comparison of cell atlases. Cell Reports 34, 108803.

Wang, S.H., Zhao, L.Y., Liu, L.S., Yang, D.W., Khatiwada, J.R., Wang, B., and Jiang, J.P. (2017). A complete embryonic developmental table of *Microhyla fissipes* (Amphibia, Anura, Microhylidae). Asian Herpetological Research 8, 108–117.

Weaver, T.E., and Conkright, J.J. (2001). Function of surfactant proteins B and C. Annual Review of Physiology 63, 555–578.

Whitsett, J.A., Kalin, T.V., Xu, Y., and Kalinichenko, V.V. (2019). Building and regenerating the lung cell by Cell. Physiological Reviews 99, 513–554.

Xu, Y., Mizuno, T., Sridharan, A., Du, Y., Guo, M., Tang, J., Wikenheiser-Brokamp, K.A., Perl, A.T., Funari, V.A., Gokey, J.J., et al. (2016). Single-cell RNA sequencing identifies diverse roles of epithelial cells in idiopathic pulmonary fibrosis. JCI Insight 1, e90558.

Yuan, B.B., Li, C.G., Kimura, S., Engelhardt, R.T., Smith, B.R., and Minoo, P. (2000). Inhibition of distal lung morphogenesis in Nkx2.1 (-/-) embryos. Developmental Dynamics 217, 180–190.

Zhang, W., Menke, D.B., Jiang, M., Chen, H., Warburton, D., Turcatel, G., Lu, C.-H., Xu, W., Luo, Y., and Shi, W. (2013). Spatial-temporal targeting of lung-specific mesenchyme by a Tbx4enhancer. BMC Biology 11, 111.

Zoetis, T., and Hurtt, M.E. (2003). Species comparison of lung development. Birth Defects Research Part B: Developmental and Reproductive Toxicology 68, 121–124.

